# A brain-wide, trial- and time-dependent deterministic drive synergizes with within-trial noise to time self-initiated actions

**DOI:** 10.1101/2025.10.28.685235

**Authors:** Michaël A. Elbaz, Kole Butterer, Sara A. Solla, Joshua I. Glaser, Andrew Miri

**Affiliations:** Department of Neurobiology, Northwestern University; Department of Human Genetics, University of Chicago; Department of Neuroscience, Northwestern University; Department of Physics and Astronomy, Northwestern University; NSF-Simons National Institute for Theory and Mathematics in Biology; Department of Neurology, Northwestern University; Department of Computer Science, Northwestern University

## Abstract

Deciding when to act in the absence of external cues is essential for exploration, learning, and survival. Yet the neural mechanisms underlying such decisions remain controversial, with current views favoring either deterministic or stochastic underpinnings. We simultaneously recorded from large neuronal populations in cortical, thalamic, pallidal, and cerebellar regions as mice self-initiated voluntary actions. Action onset timing was predictable from firing patterns up to several seconds in advance with predictions correlated across regions, demonstrating a prominent deterministic drive that spans regions. Computational modeling indicated that this drive has an initial value and rate that vary trial-by-trial, and the rate increases within trials. Although the deterministic drive is sufficient to trigger action, noise within trials also contributes to setting action timing. Therefore, discrete (across-trial) and continuous (within-trial) sources of variability synergize to time self-initiated actions. This synergy is observed brain-wide, suggesting a distributed decision-making process rather than a hierarchical, modular one.

## Introduction

The adaptive value of cognition manifests not only in deciding how to act, but also when to act. For instance, prey increase their chances of survival by fleeing at a moment predators cannot predict, rather than waiting for predators to give chase^1,2^. This raises a fundamental question: what neural mechanisms underlie action timing decisions in the absence of external cues? Seminal studies in humans over 60 years ago established the existence of the “readiness potential”, a slow buildup in electroencephalographic recordings (EEG) that precedes the onset of self-initiated actions by up to several seconds^3-5^. Correlates of the readiness potential have since been found across numerous mammalian species^6-9^, and even in fish^10^ and invertebrates^11,12^, perhaps reflecting a broadly conserved neural dynamics^13,14^.

The apparent monotonicity of the readiness potential, and classic experiments suggesting the potential begins before subjects are aware of their desire to act^4,15-18^, led to the claim that its onset reflects the decision to act, with the buildup thereafter proceeding deterministically toward action initiation^3,7,14,19-22^. However, more recent studies have challenged this classic interpretation by implicating a prominent stochastic process in setting self-initiated action timing^14,23-28^. Accordingly, the monotonicity of the readiness potential has been reinterpreted as an artifact of trial-averaging – a method commonly used with noninvasive recording techniques that yield a low signal-to-noise ratio^29^. Indeed, if action initiation requires crossing an activity threshold^7,14^, averaging neural activity aligned on action onset will produce an approximately monotonic trajectory, even if the underlying activity is dominated by random fluctuations on individual trials.

This alternative, stochastic interpretation has been formalized using drift-diffusion models (DDM), which were previously used to describe perceptual decision-making^30-36^. DDMs combine contributions from deterministic and stochastic components in a balance that, in perceptually guided tasks, varies depending on the strength of sensory evidence: the stronger the sensory evidence, the larger the deterministic component^32^. In the case of self-initiated actions, the deterministic component was assumed to be minor, insufficient on its own to reach the activity threshold beyond which an action is initiated^23,37^. Conversely, the stochastic component, thought to derive from random fluctuations in the firing rates of neurons underlying the decision^38,39^, was posited to play a prominent role, being necessary to reach the activity threshold and thereby setting action timing^14^. This view well accounts for the variable and seemingly unpredictable timing of self-initiated actions. Nevertheless, other models not involving stochastic fluctuations within trials can also account for this timing variability, such as a deterministic process that starts at random times or whose parameters vary randomly from trial to trial. Here the timing decision is made whenever a premovement buildup starts, consistent with the classic interpretation of the readiness potential. Without measurements that resolve the decision-related neural dynamics on individual trials, the predictions of these conflicting models appear equally plausible^37^, leaving the field at a conceptual impasse.

Compounding matters, the contributions of different brain regions to action timing decisions remain poorly resolved. Activity modulation preceding the onset of self-initiated actions has been observed in numerous cortical and subcortical regions^40-48^. Lesions^49-58^ and other activity perturbations^59-64^ in some regions alter action timing. Previous studies have promoted a hierarchical view, in which timing decisions rely on distinct modular contributions from different regions carried out in sequence^16,40,45-47,59,61,65,66^. In line with this, functional imaging in humans has shown a continuous sequence of activations across different cortical regions as action onset approaches^16^. However, this interpretation of hierarchical, modular contributions is based on activity measurements with low temporal resolution or that involve a limited number of neurons recorded in one or two regions at a time. Thus, it is possible that self-initiated action decisions instead rely on a shared, distributed process to which many brain regions simultaneously contribute.

To address these issues, we resolved single-unit, single-trial dynamics across large neuronal populations in eight brain regions simultaneously, in mice making self-initiated action decisions. We demonstrate that action onset timing is predictable up to several seconds prior to onset from neural activity. This predictive activity reflects a prominent deterministic drive that is temporally aligned across cortical, thalamic, pallidal, and cerebellar regions. By developing a DDM variant that accounts for the decision process at a single trial resolution, we show that the deterministic drive is parametrized differently across trials, contributing to action timing variability, and is on its own sufficient to trigger action. Yet, *in silico* ablation analyses reveal that within-trial noise also contributes to action timing and its variability across trials. The same synergistic decision process is observable across disparate brain regions. In total, our results support a hybrid model in which discrete (trial-to-trial) and continuous (within-trial) sources of timing variability act synergistically to set action timing. This resolves the prevailing dichotomy in which timing decisions were seen as either fully deterministic once a buildup process starts or primarily stochastic based on within-trial noise.

## Results

### A new behavioral paradigm to elicit self-initiated action decisions in mice

Self-initiated decisions have typically been elicited in human subjects by asking them to initiate a motor action “whenever they wish” with no experimental constraints that reinforce a specific timing^3,4,15,16^. To study such decisions in mice, we leveraged a behavioral paradigm we recently developed in which mice perform a naturalistic climbing behavior in the absence of external cues^67^. Here, head-fixed mice climb with all four limbs across handholds that extend radially from a wheel positioned in front of them, thereby rotating the wheel (Figure 1A). In the original version of the task, mice can initiate climbing at any time and are rewarded after climbing. This self-paced paradigm therefore implicitly reinforces a rapid succession of climbing bouts that maximizes the reward rate. To elicit actions whose timing is not influenced by external factors like the pursuit of rewards, we developed a variant of this task.

**Figure 1.**
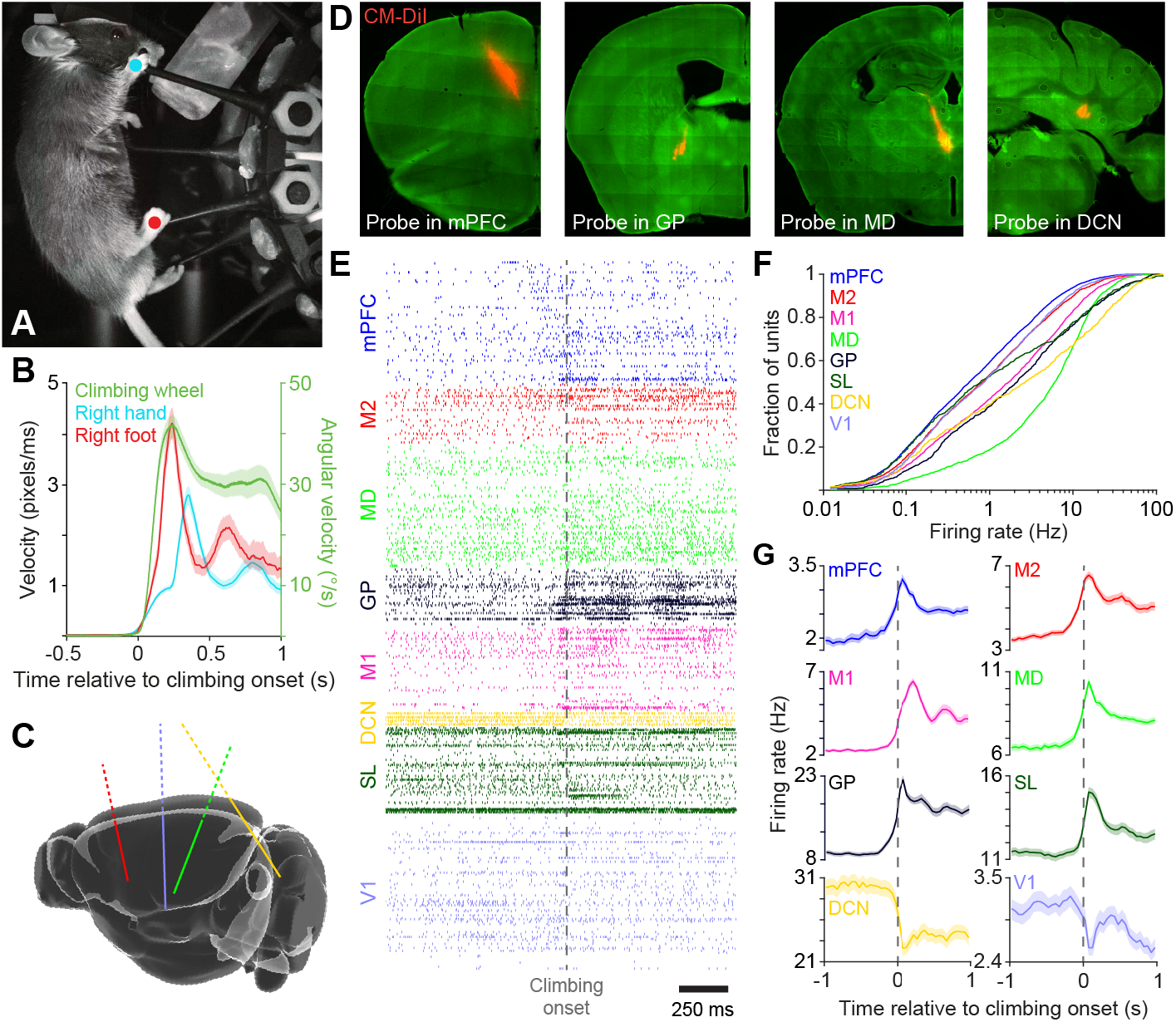
Large-scale neural recordings during self-initiated climbing. **(A)** Head-fixed mouse climbing a wheel equipped with radial handholds. **(B)** Trial-averaged velocity of the right hand and foot (left axis) and angular velocity of the climbing wheel (right axis), aligned to climbing onset (mean ± 95% confidence interval estimated from bootstrapping). **(C)** 3D schematic of a mouse brain depicting the trajectories of acutely implanted Neuropixels arrays targeting cortical, thalamic, pallidal, and cerebellar regions. **(D)** Histological sections where probe tracks appear marked with CM-DiI in the medial prefrontal cortex (mPFC), globus pallidus (GP), mediodorsal nucleus of the thalamus (MD), and deep cerebellar nuclei (DCN). Background tissue autofluorescence is shown in green. **(E)** An example spike raster surrounding climbing onset, for all eight regions of interest: mPFC, M2 (secondary motor cortex), M1 (primary motor cortex), MD, GP, DCN, SL (simple lobule) and V1 (primary visual cortex). **(F)** Empirical cumulative distribution functions of firing rates for all qualifying units in each region of interest, pooled across all recording sessions. **(G)** Trial-averaged firing rates (mean ± 95% confidence interval estimated from bootstrapping) across all qualifying units in each region, aligned to climbing onset.

Our task variant consists of alternating “GO” and “NO-GO” periods, which are distinguishable by the presence or absence of sensory cues. When no movement occurs, both periods last 3 seconds on average (exact lengths are jittered). If the mouse climbs during a GO period, a reward is delivered, the period ends and a NO-GO period begins. If the mouse climbs during a NO-GO period, the period timer is reset, delaying the start of the next GO period. Climbing onset times were computed using video-based limb tracking^68^ during epochs surrounding the onset of wheel rotation (Figure 1B). All animals trained on this task learned to climb promptly after the start of a GO period (Figures S1A-S1B). All trained animals also occasionally initiated climbing during NO-GO periods (Figure S1C), with an average of 75 ± 27 (mean ± sem) NO-GO climbing bouts per session. Analysis of anticipatory licking behavior indicated that mice are less likely to expect a reward during NO-GO periods than during GO periods (Figure S1D). This argues against the possibility that climbing bouts occurring during NO-GO periods are reward-motivated errors, wherein mice mistake a NO-GO period for a GO period. Thus, NO-GO climbing bouts do not appear reward-driven but self-initiated. We focus on these events in what follows.

### Self-initiated action timing is predictable from spiking activity

We next sought to examine spiking activity across the brain that precedes self-initiated climbing onsets. Four large-scale multielectrode arrays (Neuropixels 1.0^69^) were acutely implanted at the beginning of climbing sessions (six mice, one recording session per mouse). We targeted a range of brain regions in the left hemisphere that have previously been implicated in self-initiated action timing, or which receive direct projections from implicated regions (Figures 1C-1F, S1E and S1F, Table S1): the medial prefrontal cortex (mPFC)^46,48,59,61^, the forelimb primary motor cortex (M1)^6,8,20,70^, the forelimb secondary motor cortex (M2)^25,46^, the mediodorsal nucleus of the thalamus (MD)^71^, the globus pallidus (GP)^44,47,59,64,72,73^ and, contralaterally, the deep cerebellar nuclei (interposed and dentate nuclei, DCN) and the cerebellar simple lobule (SL)^42,43,51,62^. The array targeting cerebellar regions also recorded activity in the primary visual cortex (V1) which we included among our regions of interest, given recent evidence that this region exhibits movement-related activity^74,75^. The number of single units meeting our quality criteria (“qualifying units”) in individual regions ranged from 35 (median across sessions, DCN) to 398 (mPFC; Figure S1E). In all regions of interest, trial-averaged single unit activity appeared modulated around climbing onset and during ongoing climbing (Figure 1G).

To test for a deterministic process underlying the action timing decisions mice made, we investigated whether the timing of climbing onset could be predicted from the preceding spiking activity across regions. For each recording session, we trained a cross-validated elastic net linear regression model^76^ to predict the latency to climbing onset (time-to-movement) from the activity of qualifying units between immobility onset and climbing onset. The average duration of epochs from immobility onset to climbing onset (“trials”) ranged from 1.7 to 3.6 s across mice (Figure S1G), after ignoring here and in all subsequent analysis trials lasting less than 1 s. Firing rates were averaged in 50 ms bins, and the forecast model used the rates from the preceding five bins (250 ms) to predict time-to-movement at each time bin. Predictions for held-out trials often showed a ramping time course (Figures 2A and 2B), but were not consistent with neural activity merely tracking the passage of time (Figures S2A-S2C)^77^.

**Figure 2.**
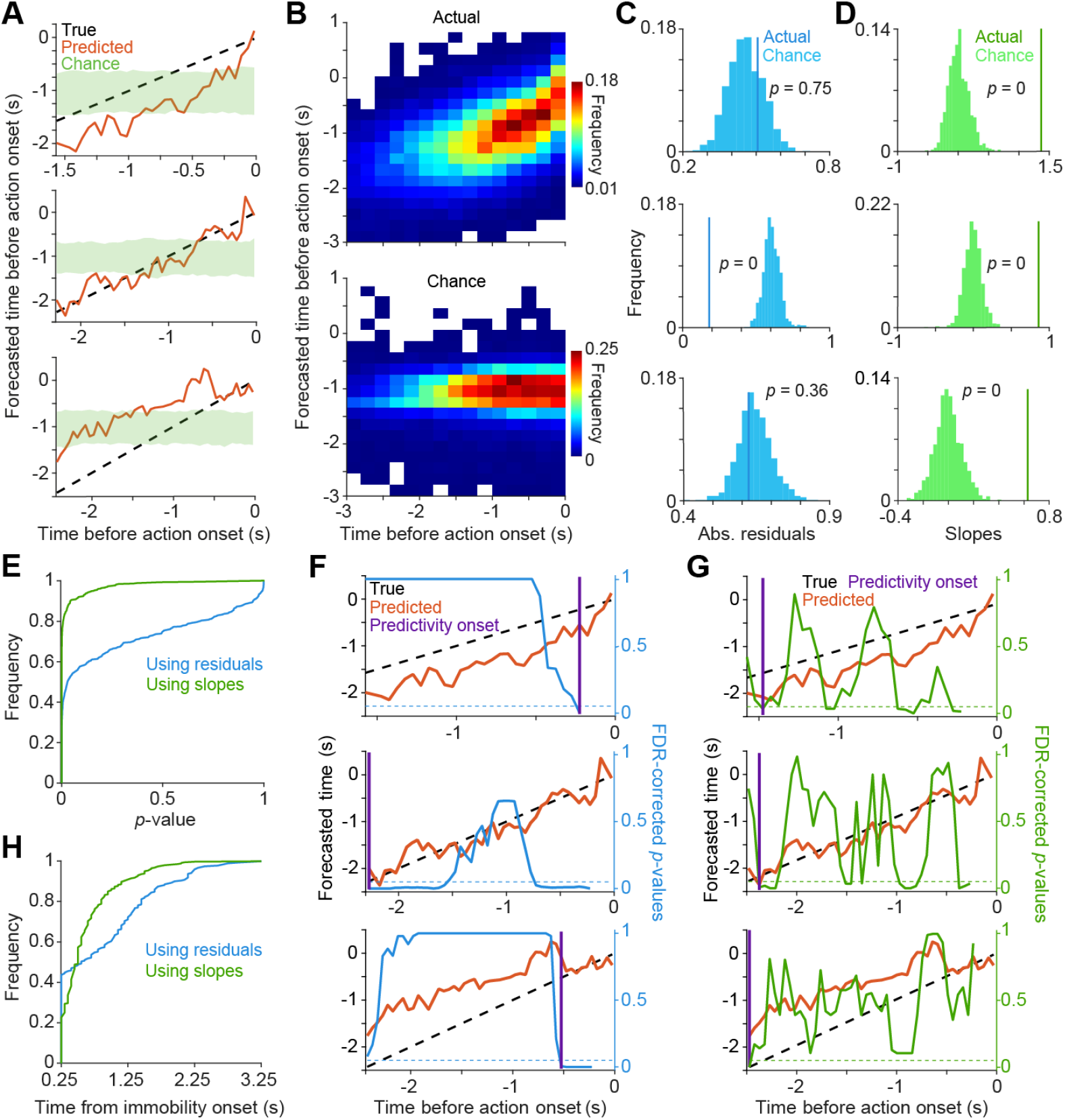
Self-initiated action timing is predictable from neural activity on individual trials. **(A)** Three example single-trial predictions of time-to-movement (red) versus true time-to-movement (dashed black). The green shaded area represents the 95% confidence interval of predictions from permuted data. Here and in subsequent analyses, all figures illustrate performance on held-out data. **(B)** 2D histograms of forecasted versus actual time-to-movement for all trials pooled across all sessions (top) and after shuffling (bottom). **(C, D)** Absolute residual (C) and slope (D) for each example trial from (A), compared with their corresponding null distributions computed with shuffling. **(E)** Empirical cumulative distribution functions of *p*-values from tests on absolute residuals (blue) and slopes (green) across all trials and sessions. **(F, G)** Identification of predictability onset for the example trials shown in (A) using either the absolute residuals (F) or slopes (G) calculated in 250-ms sliding windows. Predictability onset was defined as the first time step when the forecast traces drop below an FDR-corrected *p*-value of 0.05. **(H)** Empirical cumulative distribution functions of predictability onset times relative to immobility onset for all trials, determined using either absolute residuals (blue) or slopes (green).

To determine whether the model’s predictive accuracy was significantly better than chance, we computed two performance metrics for each held-out trial: the mean absolute difference between the forecasted and true times-to-movement (residuals), and the slope of a linear fit to the forecasted time-to-movement over time. To compute a *p*-value for each metric on each trial, actual values were compared against null distributions generated using the forecast values of models fit to permuted data. Forecast values were significantly better than chance for a majority of trials based on their residuals, and for nearly all trials based on their slopes (Figures 2C-2E). Importantly, when either performance metric was examined as a function of the time elapsed between the last GO to NO-GO period transition and immobility onset, no significant correlation was observed (Figures S2D and S2E). This argues against the possibility that models exploit lingering sensory-related activity associated with GO to NO-GO transitions. Together, these results indicate the presence of a prominent deterministic component in action timing on individual trials.

To evaluate when predictive information emerges on individual trials, we computed the same two performance metrics within 250-ms sliding windows leading up to climbing onset and compared the results to chance expectations (Figures 2F-2G). For each metric, significant predictivity emerged within the first 500 ms after immobility onset on a majority of trials (Figure 2H). This corresponds to 1.4 s ± 0.66 (median ± median absolute deviation) before action onset across all trials (Figure S2F), and to 2.78 s ± 0.58 for trials with durations ≥ 3 s (Figures S2G and S2H). This indicates that neural activity is predictive of self-initiated action timing on single trials, up to several seconds before action onset. This identification of activity predictive of action timing through our regression-based approach extends prior studies that relied on classification to predict whether an action would occur within pre-defined temporal windows^16,20^.

### Decision-related, temporally aligned activity across disparate brain regions

Is the activity used by the forecast model to predict time-to-movement localized in a subset of brain regions? To address this, we tested whether each unit’s activity improved time-to-movement predictions compared to circularly permuted controls. In all eight regions of interest, we found neurons that significantly improved forecast accuracy (“forecasting neurons”; Figure 3A), ranging from 27% (median, mPFC) to 39% (M2; Figure 3B) of units in each region. When aligned on climbing onset, the trial-averaged activity of these neurons often exhibited a ramping time course leading up to action onset (Figure 3C). This pattern was also often apparent on single trials (Figure S3A). Thus, the activity of neurons within each region of interest carries information about future action timing.

**Figure 3.**
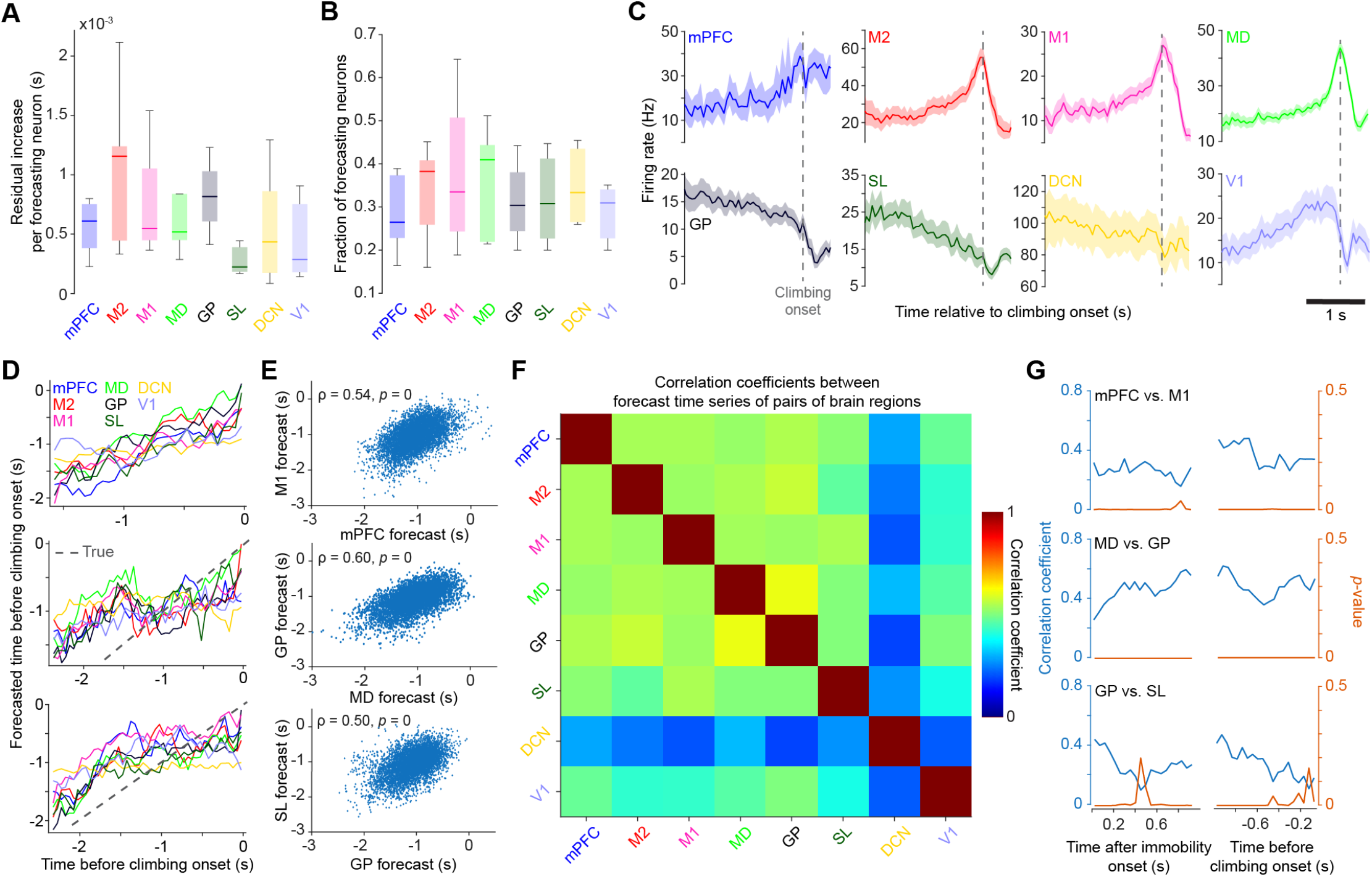
Decision-related activity is present and temporally aligned brain-wide. **(A)** Increase in forecast error upon removal of individual forecasting neurons from each region across all sessions. **(B)** Proportion of forecasting neurons in each region across all sessions. **(C)** Trial-averaged firing rate of example forecasting neurons from each of the eight regions, aligned to climbing onset (mean ± 95% confidence interval estimated from bootstrapping). **(D)** Single-region time-to-movement predictions for three example held-out single trials. **(E)** Scatterplots of time-tomovement predictions for two regions plotted against one another across all time points and trials from one session, illustrating the correlation seen between regions. ρ denotes the Pearson correlation coefficient. **(F)** Correlation matrix for single-region time-to-movement predictions. All correlations are significantly positive (Student’s t-test, FDR-corrected *p*-value < 0.05). **(G)** Crossregion correlation (blue) and associated *p*-values (red) calculated at each time step for the singleregion time-to-movement predictions of three region pairs across all trials from one session.

We then examined how decision-related activity in different regions correlates on individual trials. We trained and tested separate forecast models using activity from each individual region. The resulting single-trial forecast traces were remarkably similar across all regions, often exhibiting coordinated deviations from an ideal linear prediction of time-to-movement (Figure 3D). Predictions were positively correlated across all region pairs when pooled across all test trials and all time points up to 250 ms before action onset to avoid activity dynamics related to movement execution^15,78^ (Figures 3E and 3F). Moreover, when these across-region correlations were calculated for each 50-ms time bin, they remained consistently positive and significant throughout trials, from their emergence soon after immobility onset leading up to action onset (Figure 3G). To test whether predictive activity in one region consistently led predictive activity in another region, we computed the cross-correlations of the single-trial predictions for all region pairs. The maximum crosscorrelation peaked at or near a time lag of zero for all pairs (Figures S3B-S3E). This temporal alignment held even when predictions used firing rates in shorter, 10-ms bins and did not rely on additional preceding bins (Figures S3C and S3E). This brain-wide temporal alignment throughout the decision epoch suggests that self-initiated action timing emerges from a unified, distributed process rather than a modular, sequential one.

### The deterministic drive varies within and across trials

The preceding results demonstrate the existence of a substantial, brain-wide deterministic drive underlying self-initiated action timing, in contrast to recent models that center on the contribution of a stochastic process within trials. However, we also found that trial durations varied greatly (Figure S1G) and predictions exhibited substantial errors (Figures 2A and 2E-2G). This variability and unpredictability of action timing could reflect a substantial additional contribution from a stochastic process within trials, but could also arise from features of a deterministic process. To clarify the nature and relative prominence of the deterministic drive, we thus sought to assess different potential sources of variability in action timing.

We considered four distinct mechanisms. First, stochastic fluctuations in neural activity during individual trials (within-trial noise) might influence the decision process, as in the typical DDM. Second, timing variability could be attributed to trial-to-trial variation in the parameters of a deterministic drive. For instance, timing variability could arise from a ramping process whose initial value and slope vary across trials^37,79^. Third, the decision process might start at variable latencies after immobility onset, generating timing variability even if the subsequent dynamics are similar across trials. Finally, timing variability could also arise from trial-to-trial differences in the activity threshold.

To test these possibilities, we leveraged the output of our multi-region, time-to-movement forecast model as a single-trial readout of the decision variable, i.e., the evolution of the decision process over time. If deterministic drive parameters vary across trials, trial duration should correlate with the forecast output’s initial value and/or its slope. If the decision process starts at variable latencies after immobility onset, trial duration should be largely uncorrelated with the forecast output’s initial value. If the activity threshold varies across trials, trial duration should correlate with the forecast output’s final value.

Thus we first plotted three variables versus trial duration: the forecast output’s initial value (250 ms after immobility onset because of the window used for prediction), its slope when best fit with a line, and its final value (250 before action onset or at action onset). We found that the initial values and the slopes were negatively correlated with trial duration in all six mice (Figures 4A and 4B). By contrast, the final values, though highly variable, were uncorrelated with trial duration (Figures 4C and 4D). To ensure that the negative correlation between initial value and trial duration was robust across trials, and not due to a subset of trials with an exceptionally early decision process start time, we confirmed that the negative correlation holds for subsets of trials (Figure S4). This also further supports an early engagement of the decision process soon after immobility onset. Together, these results suggest that variability in action timing stems from variation in deterministic drive parameters across trials, but not in the start time or activity threshold of the decision process.

**Figure 4.**
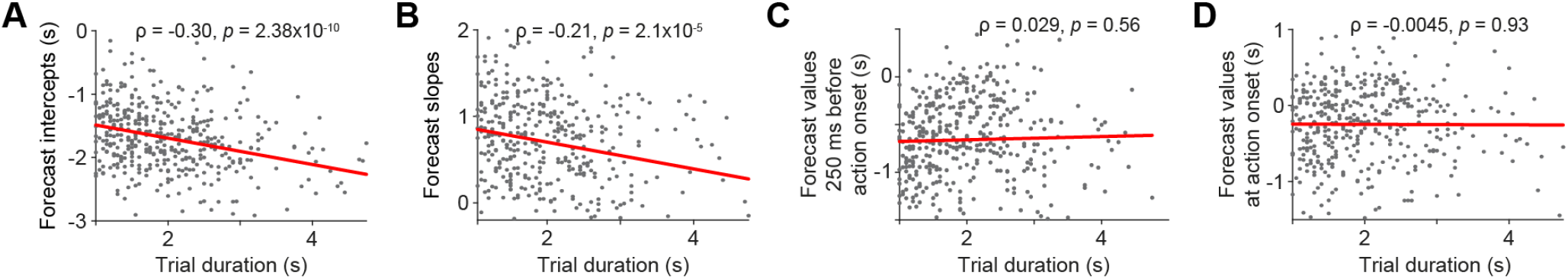
Action timing correlates with the initial value and slope of the forecast model output, but not its final value. **(A-D)** Scatterplots of the initial value (A), slope of a linear fit (B), values 250 ms prior to climbing onset (C), and final value at climbing onset (D) for forecast traces versus trial duration across held-out trials from all sessions. ρ denotes the Pearson correlation coefficient, with *p-*values determined by a Student’s t-test. Red lines show a best-fit line.

To further probe for trial-to-trial variation in the parameters of a deterministic drive, we used model selection. In the typical DDM, the decision variable *x* has an initial value *x*_0_, and three other parameters govern Δ*x*_*n*_, the change in the decision variable at time *t*_*n*_ [Equation 1]: *I* is the drift rate, the slope of a linear ramp; *k* is the amplitude of a leak process that acts as a restoring force towards baseline^33^; and *c* is the amplitude of a diffusion process that models the accumulation of stochastic fluctuations within trials, with *ξ*_*n*_ drawn from a standard normal distribution. Thus, in this model, the approach of the decision variable to the activity threshold effectively depends on deterministic drift and leak processes, as well as the accumulation of within-trial noise.

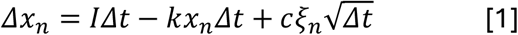

In this model, since the values of the four parameters are constant across trials, timing variability across trials can arise only from the diffusion process. To capture potential trial-to-trial variability in deterministic drive parameters, we formulated a DDM variant [Equations 2-5]. Here, the initial value and drift rate are drawn from normal and log-normal distributions, respectively, on individual trials. Additionally, previous studies of stimulus-driven perceptual decisions have described an apparent increase over time in drift rate within individual trials^80-88^. Thus, we also incorporated either a linear or exponential factor into the drift term to capture potential time-dependent changes in the drift rate^82,86,87^. This yielded a model with seven parameters: *μ*_*intercept*_, *σ*_*intercept*_, *μ*_*drift*_, *σ*_*drift*_, *α, k* and *c*.

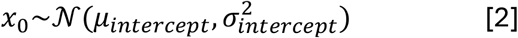

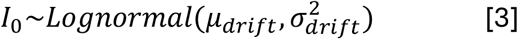

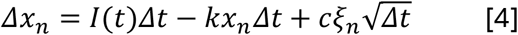

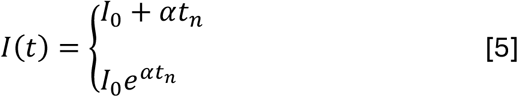

We then tested whether the inclusion of individual parameters improved fits to the forecast output. Specifically, we used maximum likelihood estimation to quantify the fit improvements on held-out forecast traces from including *σ*_*intercept*_, *μ*_*drift*_, *σ*_*drift*_, *α* and *k* (Figure S5). All models included *μ*_*intercept*_ since zero is not the average but the lowest possible initial value of our fitted forecast traces (see Methods). Our model selection analysis revealed that each tested parameter significantly improved the fit quality for each mouse (Figures 5A, 5B, S6A and S6B). This further supports the presence of trial-to-trial variability in the deterministic drive parameters (*σ*_*intercept*_ and *σ*_*drift*_).

**Figure 5.**
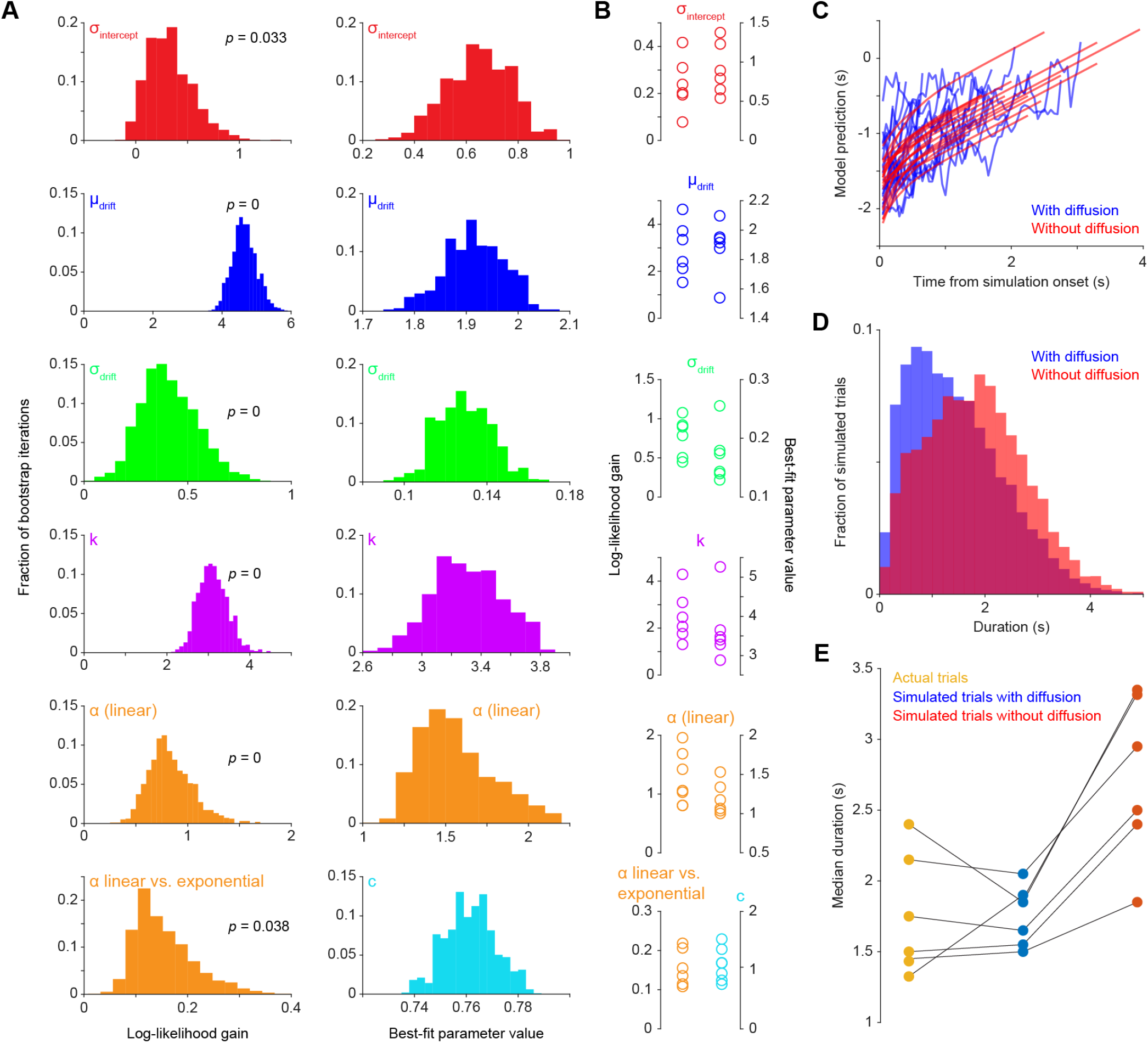
Modeling reveals a trial- and time-dependent deterministic drive and trial-shortening within-trial noise. **(A)** *Top five rows*: Histograms of the log-likelihood gain from including each parameter (rows) in the model (left column) and the best-fit parameter value (right column) across bootstrap iterations for one example session. *Bottom row*: for the same session, log-likelihood gain for linear versus exponential drift rate growth (left) and the best-fit values for the diffusion amplitude *c* (right) across bootstrap iterations. Note that *μ*_*drift*_ and *σ*_*drift*_ correspond to the mean and standard deviation of the underlying normal distribution for log(*I*_0_) (see Figures S6A and S6B for conversion to the lognormal distribution scale, and Methods). **(B)** Median log-likelihood gain (left column) and best-fit parameter values (right column) for individual mice (circles). **(C)** Simulations of the decision variable over time using the best-fit DDM for one example session, either with (blue) or after excluding (red) the diffusion term. **(D)** Distribution of simulated trials’ duration (time to threshold) using the best-fit DDM for one example session, either with (blue) or after excluding (red) the diffusion term. **(E)** Median duration of actual trials (yellow), and simulated trials using the best-fit DDM for each session, either with (blue) or after excluding (red) the diffusion term. The duration of actual trials excludes 0.2 s after immobility onset and 0.25 s before action onset as the corresponding time-to-movement predictions, which are fit to the DDMs, exclude these epochs (Methods). Accordingly, the first 0.2 s of simulated trials are discarded before computing their duration.

In addition, we found that best-fit *α* values were positive, indicating that the drift rate increases over time within a trial, with a greater likelihood gain achieved with a linear rather than an exponential growth (Figures 5A and 5B). The best-fit values of *α* imply a ∼1.5-fold increase for a 2-s trial, indicating a substantial contribution to action timing from this factor. Thus, the deterministic drive effectively increases in strength as time passes, like an increasing urgency signal.

As a control, we assessed whether variability in the initial value (*σ*_*intercept*_) might actually reflect within-trial noise accumulated prior to immobility onset, during the execution of the previous movement^29^. We calculated the theoretical variance limit of the system (*c*^2^/2*k*), which represents the equilibrium between noise accumulation and leak decay^89,90^. We found that the observed initial variance 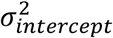 consistently and substantially exceeded this limit (Figure S6C). This implies that *σ*_*intercept*_ is not a byproduct of pre-trial noise accumulation, but instead reflects a state at immobility onset that varies across trials.

Altogether, these observations are consistent with the decision process involving a deterministic drive that varies (a) discretely from trial to trial in initial value and rate, and (b) over time within a trial, having a drift rate that increases over time.

### Within-trial noise is not required for action initiation but contributes to action timing

Having established the trial- and time-dependent nature of the deterministic drive, we then sought to examine how within-trial noise contributes to action timing. Previous work using the typical DDM to account for trial-averaged EEG data has suggested that within-trial noise plays a prominent role, being necessary to reach the activity threshold (Figure S6D)^23,37^. Since noisy fluctuations are inherent to neural activity and, consequently, to our forecast traces, a diffusion term will necessarily improve the fit of our models to forecast model output. Thus we did not test whether including the diffusion term improved fits, but rather investigated its contribution to timing variability and whether it is necessary to reach the activity threshold.

To do this, we performed an *in silico* ablation analysis: we simulated trials using our DDM variant fit to data from individual sessions, and then repeated these simulations without the diffusion term. We found that the activity threshold was reached in both cases (Figure 5C). However, including the diffusion term did shorten the duration of the trials by 22 to 45% across mice (Figures 5D and 5E). As expected, the presence of the diffusion term also increased the variance in trial duration, by 16 to 36% across mice. These results suggest that while not required to self-initiate actions, within-trial noise contributes substantially to setting action timing.

As a control, we assessed the possibility that autocorrelation in the within-trial noise may further improve our model^38,91,92^. We augmented our DDM variant with an eighth parameter, a spectral exponent *β*, which introduces dependence across time points in the diffusion term. However, adding this parameter did not yield a significantly better fit compared to using temporally uncorrelated noise (Figures S6E and S6F).

### Trial-dependent deterministic drive and trial-shortening within-trial noise are found brain-wide

Finally, we asked whether our modeling results using multi-region activity were also locally preserved within individual brain areas. We repeated the model selection with our DDM variant fit to forecast output derived from each region individually. This analysis revealed that for all eight regions of interest, model fits were significantly improved by incorporating trial-to-trial variability in the decision variable’s initial value (σ_*intercept*_) and drift rate (σ_*drift*_), as well as a within-trial increase of the drift rate (α; Figure 6A). Moreover, the best-fit parameter values were remarkably similar across regions (Figures 6B, S6G and S6H). We again found that the diffusion term hastened action onset in each region (Figure 6C). These results demonstrate that a common decision process is reflected across all regions we examined. This further supports a broadly distributed decision process coordinated across regions.

**Figure 6.**
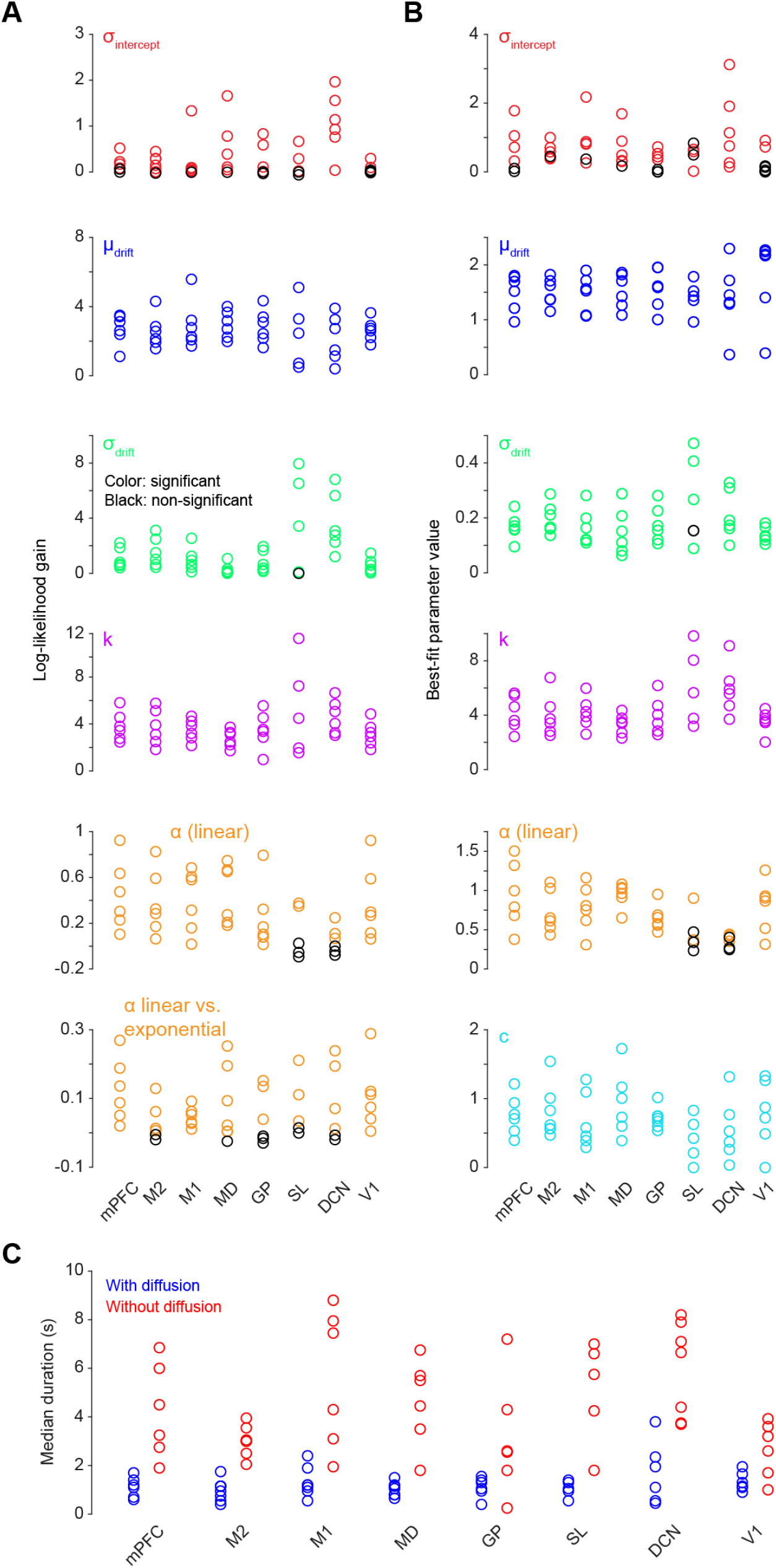
A similar decision process is reflected across brain regions. **(A)** For individual regions (columns) in each mouse (circles), the median log-likelihood gain across bootstrap iterations from including each parameter (rows) in the model using the forecast traces (top five rows) and from linear versus exponential drift rate growth (bottom row). Colored circles in (A) and (B) indicate a significant improvement in model fit. **(B)** For individual regions (columns) in each mouse (circles), the median best-fit parameter values across bootstrap iterations. Note that *μ*_*drift*_ and *σ*_*drift*_ correspond to the mean and standard deviation of the underlying normal distribution for log(*I*_0_) (see Figures S6G for conversion to the log-normal distribution scale, and Methods). **(C)** Median duration of simulations (time to threshold crossing) generated with the single-region best-fit DDMs, with (blue) and after excluding (red) the diffusion term, across sessions (circles).

## Discussion

In this work, we sought to shed light on the neural basis of self-initiated timing decisions by resolving single-trial, single-unit activity across multiple brain regions simultaneously. We found that action onset timing can be predicted from spiking activity up to several seconds in advance, and predictions are temporally aligned throughout the decision epoch across cortex, thalamus, basal ganglia, and cerebellum. This implies that a prominent, brain-wide deterministic drive underlies such decisions.

By assessing different potential sources of variability in action timing, we found that the deterministic drive varies in both its initial value and its rate across trials, and its rate increases over time within trials. In contrast to recent models, within-trial noise is not required for crossing the activity threshold, but does shorten the average time to threshold crossing. Similar decision-related activity dynamics were observed in all brain regions we examined, indicating a broadly shared decision process. Altogether, our results offer a unifying account in which trial-by-trial variability in deterministic drive parameters and within-trial noise act in concert to render the timing of self-initiated actions unpredictable.

Our results imply that a buildup of neural activity preceding action onset is a genuine feature of the decision process, not an artifact of trial-averaging. Yet, they stand in contrast to classic views of a purely deterministic process in which action onset is set whenever the buildup starts^3,4,7,14-17,19-22^. Our results also stand in contrast to more recent accounts whereby noisy neural activity within trials is the sole driver^23,28^, or principal driver^25^, of timing decisions. This aligns with a previous proposal^37^ that the early predictability of action timing shown in reference^25^ is more consistent with a strong deterministic drive than with accumulation of within-trial noise.

Our ability to resolve a long-standing debate regarding the deterministic versus stochastic underpinnings of self-initiated timing decisions stems from two key features of our approach. First, the vast majority of neural data recorded in relation to self-initiated decisions has been obtained in humans, using methods whose resolution necessitate trial-averaging. Yet single-trial resolution is required to arbitrate between competing models. For instance, in the “stochastic accumulator model”, developed by fitting the typical DDM to such trial-averaged data, within-trial noise is both the sole source of timing variability across trials and is required for threshold crossing (Figure S6D)^14,23^. However, these same data are equally well fit by models such as the linear ballistic accumulator, which lacks within-trial noise, and in which the decision process is fully deterministic once its parameters are set when the pre-movement buildup starts^37,79^. Second, previous models of self-initiated decisions have treated within-trial noise and trial-to-trial variation in deterministic drive parameters as mutually exclusive^14,37^, though the same has not been true for models of perceptual decisions^30,36^. By combining these potential mechanisms within a single model, we resolved a dichotomy and established that they act synergistically.

Here we uncovered a temporal alignment in decision-related activity across eight brain areas, from soon after immobility onset to action onset. This alignment was best illustrated by the striking similarity of the errors in time-to-movement predictions (Figure 3D). Key to this observation was our simultaneous spike-resolution recording of large neuronal populations in multiple regions, which allowed the comparison of predictive dynamics across different regions on the same single trials. Such brain-wide coordination appears to contrast with previous suggestions of a more modular organization whereby, for example, deterministic and stochastic components originate in mPFC and M2 respectively, with activities predictive of action timing confined to M2^46^. A more distributed picture emerges from our results, with predictive activity detectable in both of these regions, as well as several others. Contrary to the classical view of a hierarchical organization in which regions perform distinct roles in a feedforward sequence, our results are more consistent with a unified decision-making process that unfolds synchronously across disparate brain regions, relying on recurrent interactions between regions. These brain-wide dynamics we found in relation to self-initiated decisions echo the broad coordination reported recently during motor execution^74,75,93,94^ and perceptual decision-making^95-101^, including regions historically seen as inherently motor (e.g., M1) or sensory (e.g., V1).

Our results also reveal additional parallels with perceptual decisions, which remain far more extensively studied than self-initiated decisions. In tasks that require waiting, neural activity that builds up at rates that scale with the waiting time has been observed across cortical regions and the basal ganglia^30,73,102-106^. Moreover, our findings of a drift rate that increases within a trial are reminiscent of urgency signals proposed to underlie perceptual decisions^36,80-88^. These urgency signals have been modeled by either a time-varying drift rate or a collapsing threshold. Thus, urgency may be a general organizing principle of decision dynamics, even in the absence of reward or other incentives to act quickly. This mechanistic similarity between sensory-guided and self-initiated decisions appears well-suited for negotiating complex natural environments, where behavior is rarely dictated purely by internal or external factors, but rather emerges from a dynamic integration of both.

However, while mechanistic features appear conserved between self-initiated and sensory-guided decisions, the nature of the decision process differs fundamentally in its input and adaptive function. In perceptual tasks, the deterministic drive reflects the strength of external sensory evidence, and stochasticity primarily represents perceptual noise that can be detrimental to choice accuracy^107^. In contrast, in the absence of external cues, the deterministic drive we observed likely reflects an internal imperative to act^14^ ensuring that an action will be executed, while stochasticity gives rise to an advantageous unpredictability of behavior.

An important question unaddressed by our results concerns the neurophysiological underpinnings of the sources of action timing variability we have described. Our analysis implicates three such sources: trial-to-trial variability in the initial value and drift rate of the deterministic drive, and noise within trials. While within-trial noise is likely attributable to aspects of neuronal physiology, like ion channel gating and synaptic transmission^39^, the origins of discrete sources of variability are more obscure. Neuromodulatory systems may be one possible source. Notably, the basal forebrain cholinergic system has been causally implicated in self-initiated action timing^47,59,60^ and serves as the primary source of acetylcholine to the cortex^108^, including the prefrontal and somatomotor regions where we observed predictive activity. Cholinergic input can modulate correlations across a neuronal population^109^, and acetylcholine appears capable of modulating cortical function on the timescale of individual trials^110-116^. Thus, the basal forebrain cholinergic system could participate in regulating the gain of the deterministic drive (i.e., the drift rate) on individual trials.

## Supplementary Figures

**Figure S1.**
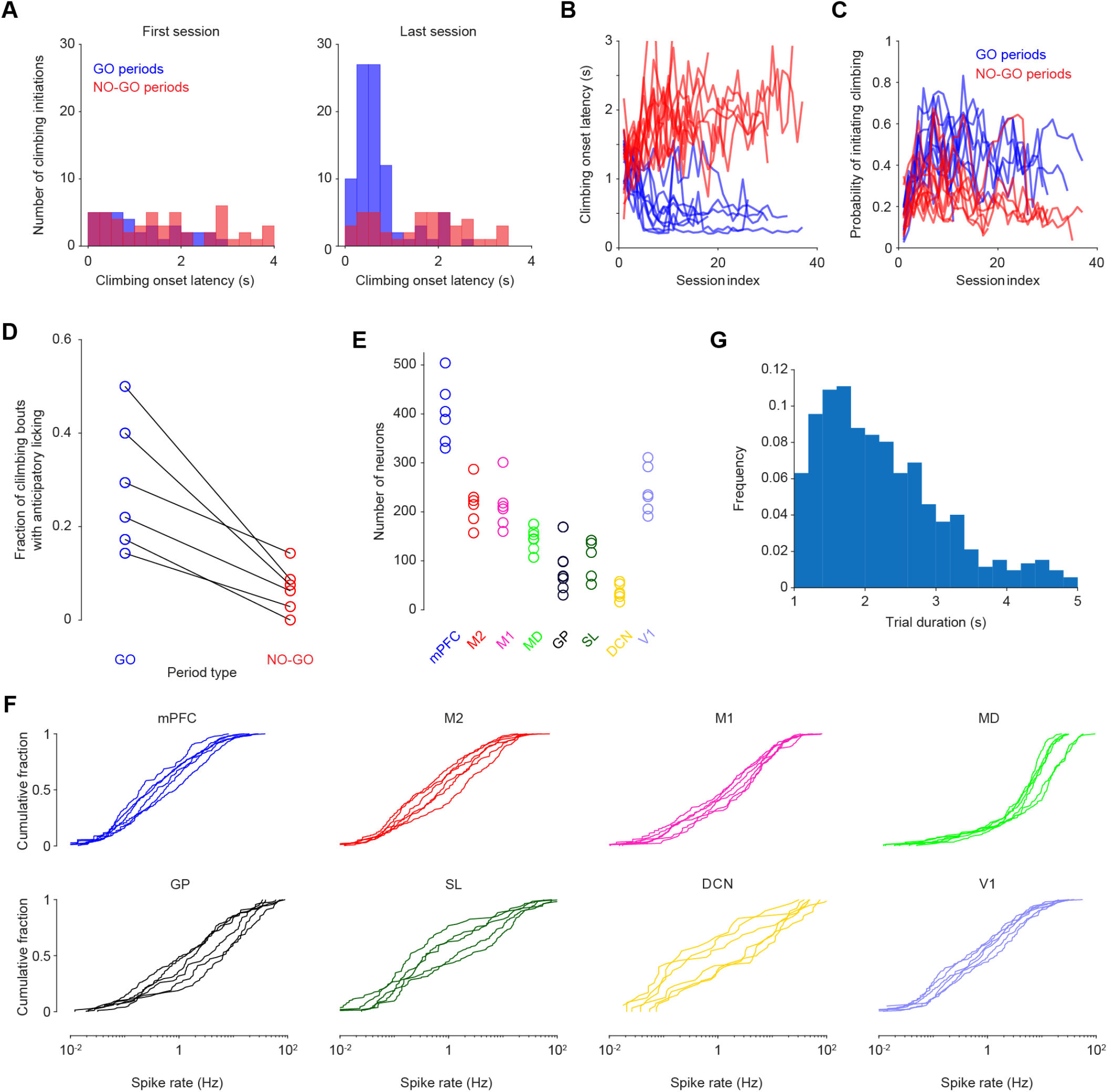
Behavioral characterization and electrophysiological yield. **(A-C)** Evolution of climbing behavior across learning. Over the course of training, animals learn to rapidly initiate climbing upon GO onset, while maintaining a baseline rate of self-initiated climbing during NO-GO periods. These panels show data from all trained animals (besides the ones for which we carried out electrophysiological recordings) across all learning sessions. Because video monitoring was only performed during electrophysiological recording sessions, climbing onset times were here determined exclusively using wheel data. For NO-GO periods, the latency of the first climbing initiation only is considered in the event of multiple initiations. (A) Histograms of climbing onset latency during GO (blue) and NO-GO (red) periods for a single representative mouse, for the first training session (left) and the last training session (right). (B) Median climbing onset latency across training sessions for all mice. Each line represents an individual mouse for GO (blue) and NO-GO (red) periods. (C) Probability of initiating climbing across training sessions for GO (blue) and NO-GO (red) periods. Each line represents an individual mouse. **(D)** Fraction of climbing bouts that included anticipatory licking during GO and NO-GO periods. This analysis is restricted to the recording sessions, where video data was recorded and used to track licking behavior. Each connected line represents data from a single recording session. **(E)** Number of qualifying single units per recorded region across all six recording sessions, with each circle representing one session. **(F)** Cumulative distributions of spike rates for single units within each region, with each trace representing one session. **(G)** Histogram of trial durations pooled across all recording sessions.

**Figure S2.**
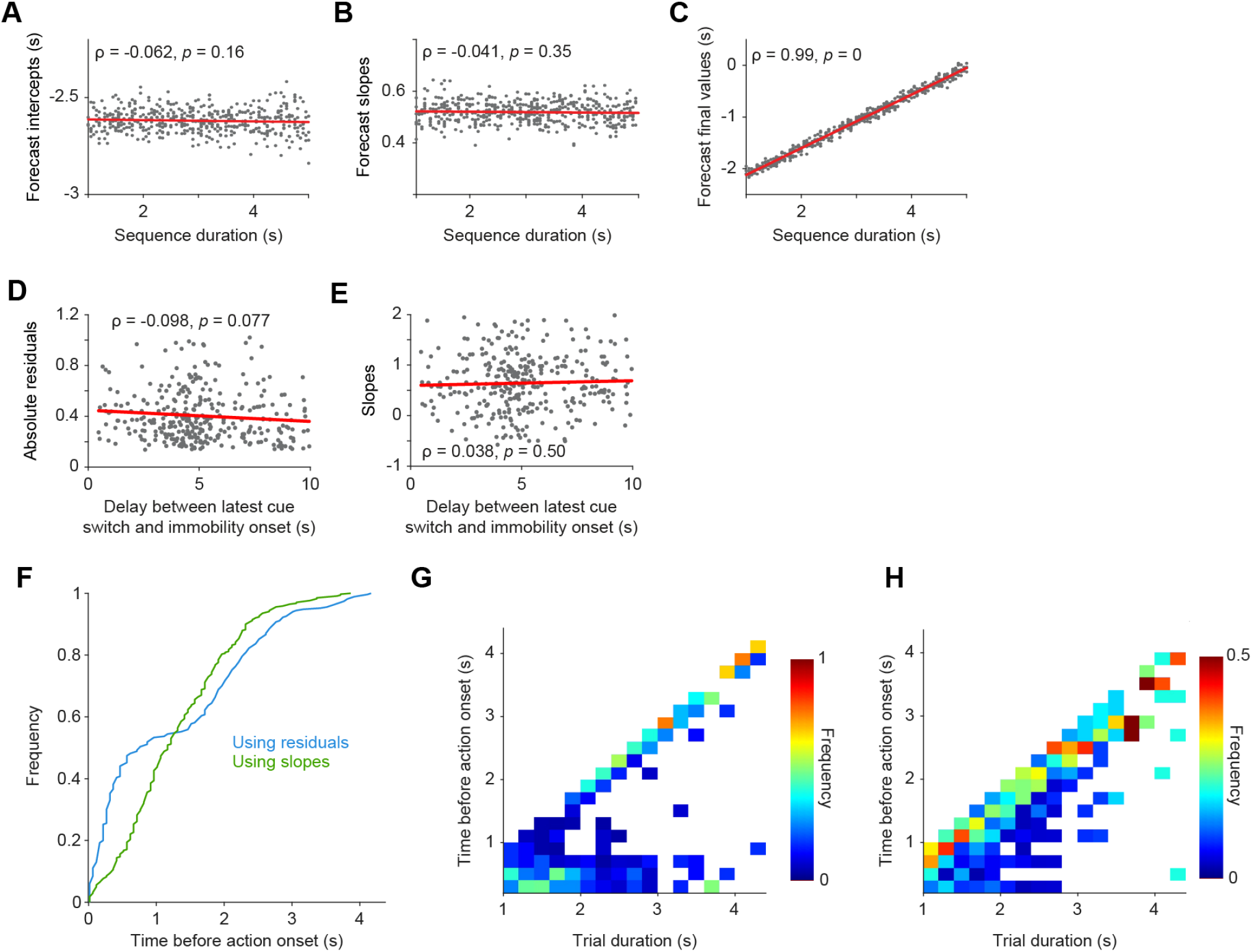
Forecast controls and emergence of single-trial predictability. **(A-C)** Properties of forecast traces obtained using synthetic “clock-like” activity. (A) Lack of correlation between the initial forecast values (intercepts) and synthetic trial duration. ρ denotes the Pearson correlation coefficient. (B) Lack of correlation between the forecast slopes and synthetic trial duration. (C) Positive correlation between the final forecast values and synthetic trial duration. These properties are qualitatively opposite to those obtained using the actual neural data (Figures 4A-4D). **(D, E)** Lack of correlation between the delay from the latest cue switch to immobility onset and either the absolute residuals (D) or slopes (E) of the forecast traces. For each trial, metrics were computed over the first second of the forecast traces. All data shown are from cross-validated, heldout test trials. **(F)** Cumulative distribution of predictability onset times relative to climbing onset across all trials. Figure 2H shows the same data relative to immobility onset. **(G, H)** 2D histograms depicting the emergence of predictability relative to climbing onset as a function of trial duration. The time of predictability emergence is calculated using either residuals (G) or slopes (H).

**Figure S3.**
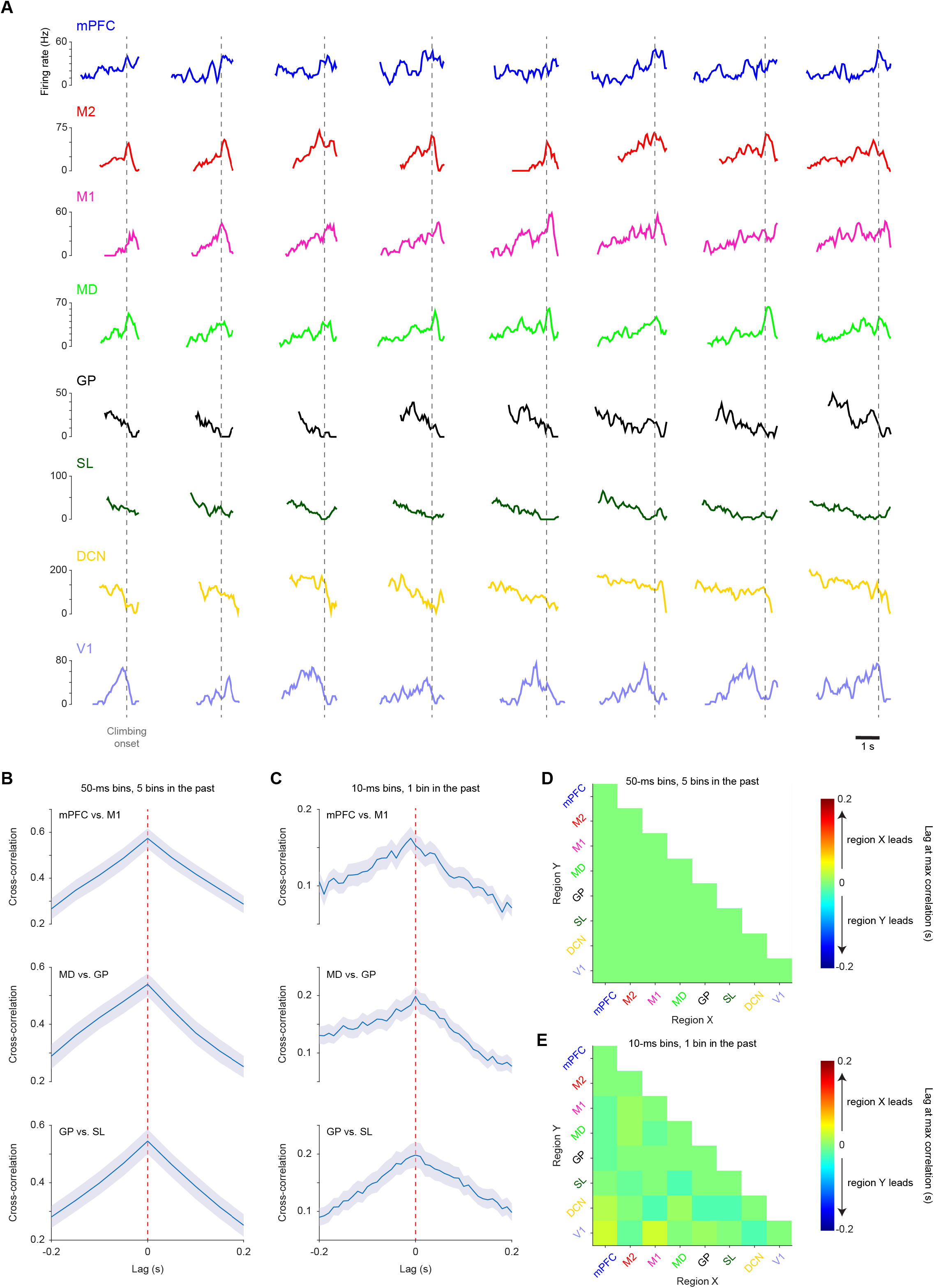
Predictive activity across brain regions. **(A)** Single-trial firing rate of an example forecasting neuron from each of the eight regions of interest, aligned to climbing onset. Causal moving average was applied: the value at each time point was calculated by averaging the data from that point with those of the preceding 200 ms window. Trials are sorted by increasing immobility durations. These example neurons correspond to the ones whose trial-averaged firing rates are shown in Figure 3C. **(B-C)** Cross-correlation of time-to-movement predictions across brain regions, averaged across trials (mean ± 95% confidence interval estimated from bootstrapping). A positive lag at maximum cross-correlation indicates that the first region listed (i.e., region X in “region X vs. region Y”) is leading. (B) Cross-correlation for three example region pairs using our standard forecast models: trained and tested with spike trains convolved with a Gaussian kernel (*σ* = 10 ms), binned into 50-ms intervals, and using 5 past bins for each prediction. (C) Crosscorrelation for the same three region pairs using forecast models trained and tested with spike trains without convolution, binned into 10-ms intervals, using 1 past bin for each prediction. **(D)** Matrix of lags at maximum cross-correlation for all region pairs using the standard forecast models described in B. **(E)** Corresponding lag matrix using the forecast models without convolution described in C. No lag was significantly different from 0 (non-parametric bootstrap test, FDR-corrected *p*-value > 0.05).

**Figure S4.**
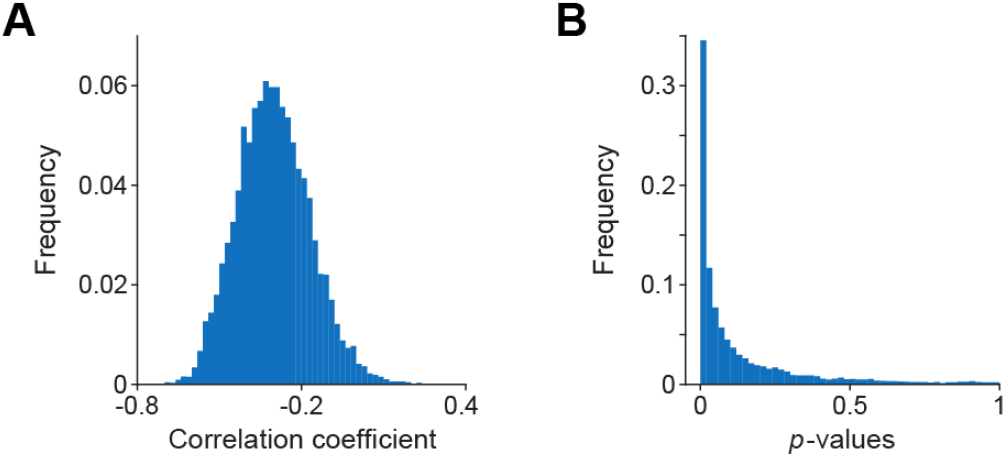
Robustness of the negative correlation between initial forecast values and trial duration (established in Figure 4A) assessed via random subsampling. **(A)** Distribution of Pearson correlation coefficients across 10,000 iterations, where 10% of the trials were randomly retained, without replacement, in each iteration. The resulting distribution remains predominantly negative and centered around the mean value obtained from the full dataset. **(B)** The corresponding distribution of *p*-values. Together, these distributions confirm the stability of the relationship even with substantially reduced statistical power.

**Figure S5.**
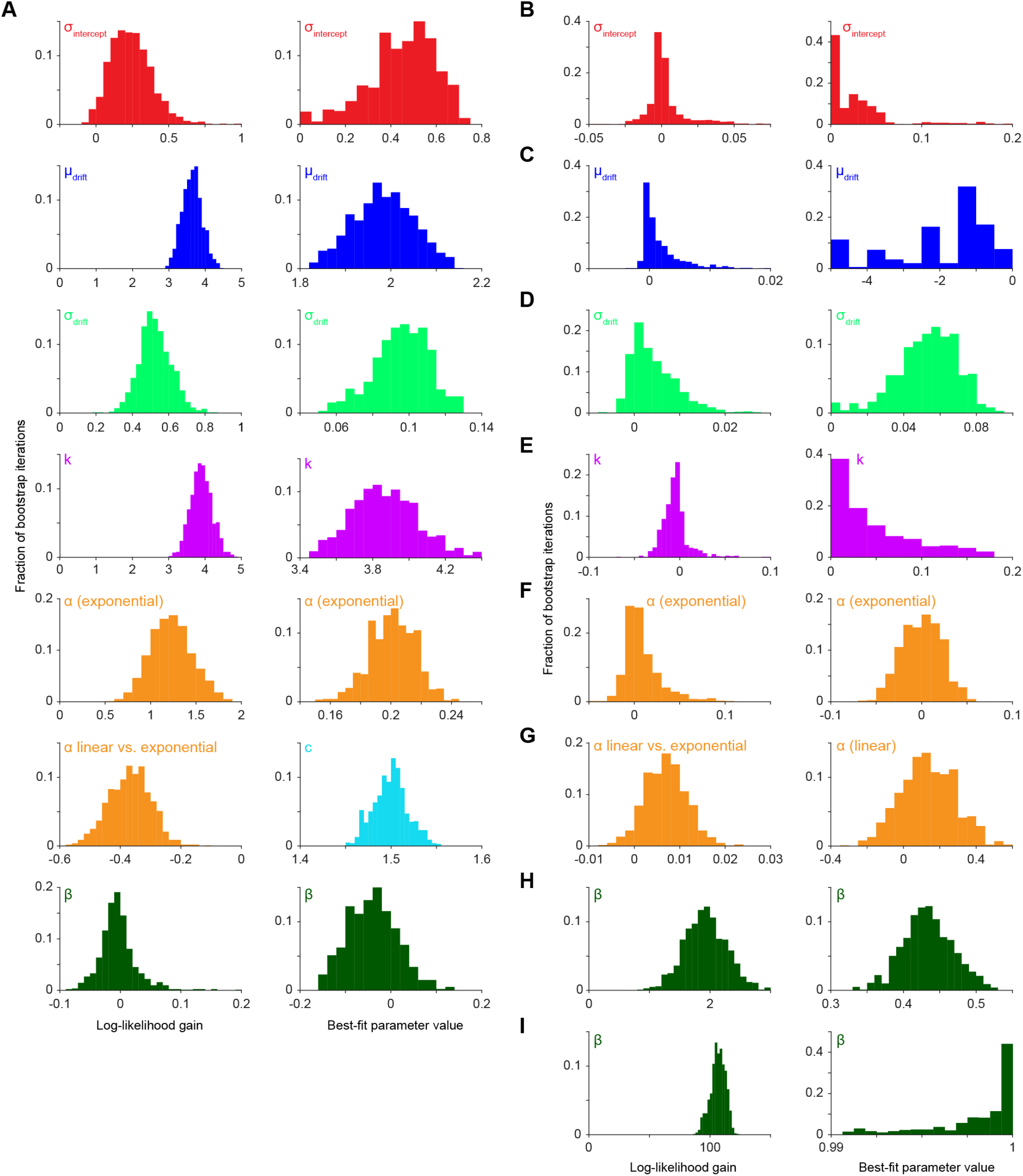
Validation of the DDM fitting and model selection procedure via parameter recovery. To mimic the fitting conditions of forecast traces, whose first point corresponds to 250 ms after immobility onset, all simulated sequences were fit after truncating their first 200 ms. Distributions show the results across bootstrap iterations. For each panel, the left column shows the loglikelihood gain, and the right column shows the best-fit parameter value. A positive likelihood value for the “*α* linear vs. exponential” comparison denotes a higher likelihood for the model with a linear growth rate. **(A)** Recovered parameter distributions for sequences generated using our 7-parameter DDM variant (*β* = 0). Ground-truth parameters for generating the sequences were: diffusion *c* = 1.5, leak *k* = 4, time-dependent drift *α* = 0.2 (exponential growth), mean initial value *μ*_*intercept*_ = 1, trial-to-trial variability in initial value *σ*_*intercept*_ = 0.5, mean drift rate *μ*_*drift*_ = 2, and trial-to-trial variability in drift rate *σ*_*drift*_ = 0.1. The procedure accurately recovers the ground-truth values for the 7 parameters and correctly identifies *β* as non-significant (log-likelihood gain ≈ 0). **(B-F)** Parameter recovery using reduced 6-parameter generative models. Sequences were generated with one specific parameter omitted (set to 0) to confirm that the model selection procedure correctly assigns a log-likelihood gain near zero for the missing parameter: (B) *σ*_*intercept*_ = 0 (similar initial value across trials), (C) *μ*_*drift*_ = 0 (no drift), (D) *σ*_*drift*_ = 0 (similar drift across trials), (E) *k* = 0 (no leak), and (F) *α* = 0 (constant drift within a trial). **(G)** Parameter recovery for sequences generated with a linear rather than exponential drift growth (*α* = 0.2). **(H, I)** Recovered parameter distributions for sequences generated with a full 8-parameter model, demonstrating the procedure’s ability to successfully extract the additional parameter *β* when set to (H) *β* = 0.5 and (I) *β* = 1.

**Figure S6.**
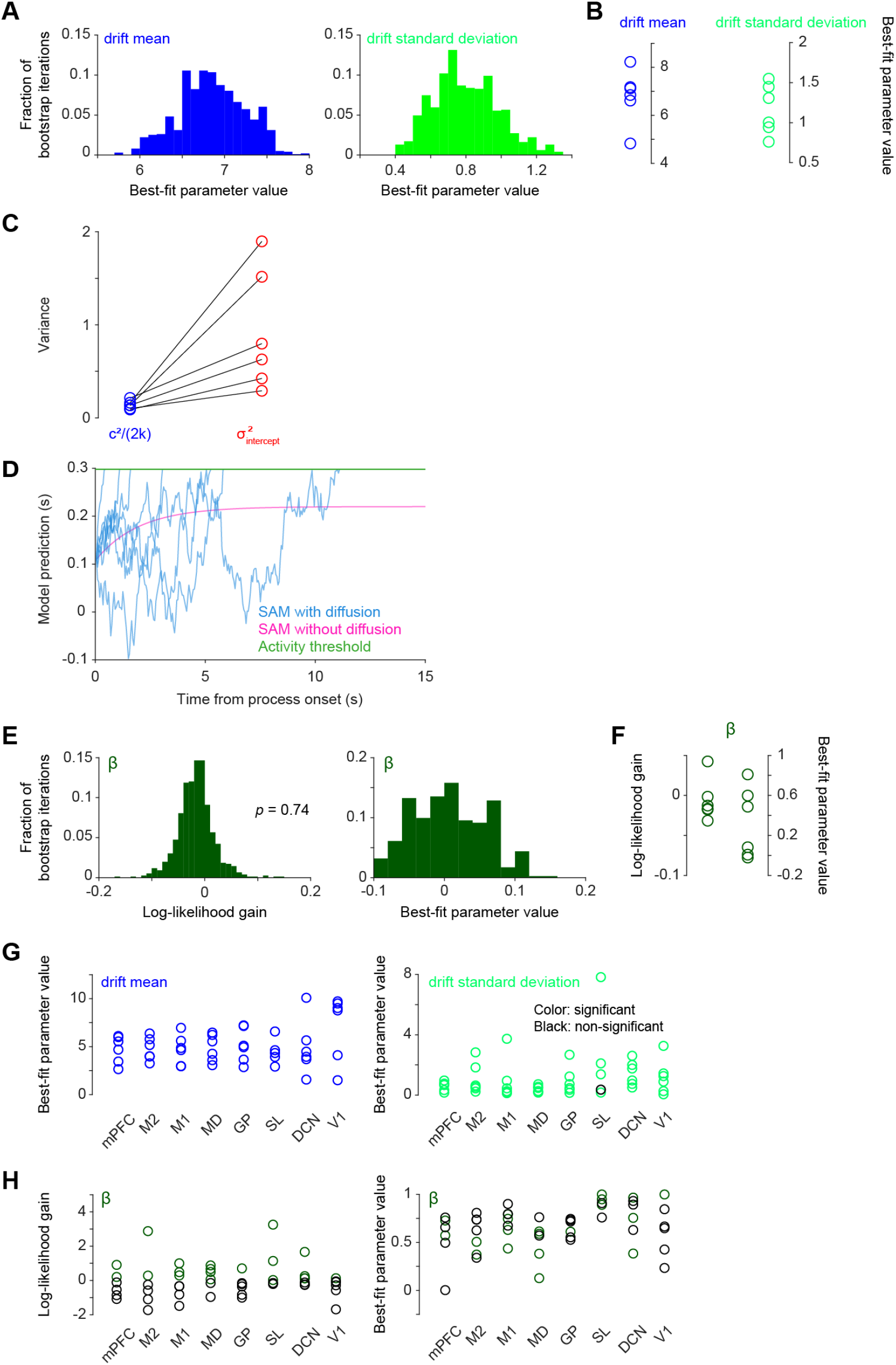
DDM controls and comparison with the “stochastic accumulator model”. **(A, B)** Best-fit parameter values for the mean and standard deviation of the initial drift rate *I*_0_. As noted in Figure 5, the fitted parameters *μ*_*drift*_ and *σ*_*drift*_ define the mean and standard deviation of the underlying normal distribution for log(*I*_0_) [Equation 3]. Here, drift mean (blue) and drift standard deviation (green) represent the corresponding mean and standard deviation of the log-normal distribution itself, from which the initial drift *I*_0_ is drawn on a per-trial basis. (A) Parameter distributions across bootstrap iterations for the same illustrative session as in Figure 5A. (B) Median best-fit parameter values for each of the six mice (corresponding to the data in Figure 5B). Each circle represents a mouse. **(C)** Comparison of steady-state variance and initial variance. Comparison between the theoretical asymptotic variance limit of the leak process (*c*^2^/(2*k*), blue) and the trial-to-trial variance in initial value (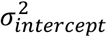, red) for each of the six mice. Lines connect data from the same recording session. **(D)** Simulations based on a typical DDM [Equation 1] using parameters from a previous study^23^ which define the “stochastic accumulator model” (SAM; activity threshold = 0.298, *I* = 0.11, *c* = 0.1, *k* = 0.5). The deterministic component alone (SAM without diffusion, magenta trace) plateaus and fails to reach the activity threshold (green line). In this framework, the addition of a diffusion term (SAM with diffusion, blue trace) is strictly required for threshold crossing. **(E, F)** Model selection provides no evidence for temporally autocorrelated (colored) noise. To test whether the diffusion term had temporal correlations, the baseline DDM was augmented with a spectral exponent *β*, where *β* = 0 corresponds to white noise. (E) Distributions of the log-likelihood gain (left) and the best-fit *β* value (right) for the same illustrative session as in Figure 5A. The inclusion of *β* did not yield a significant gain. (F) Summary of colored noise testing for all six mice. Shown are the median log-likelihood gains (left) and best-fit parameter values (right). The inclusion of *β* did not significantly improve the fit for any of the mice. Together, these results justify the use of our 7-parameter model with white noise in the Main text. **(G, H)** Region-specific parameter conversions and colored noise controls. (G) Best-fit parameter values for the mean (left) and standard deviation (right) of the initial drift rate *I*_0_ for each individual brain region across all sessions, corresponding to the models evaluated in Figure 6. (H) Region-specific evaluation of temporally autocorrelated noise (*β*). The inclusion of *β* did not significantly improve the fit for the majority of individual sessions across regions.

## Material and Methods

### Experimental subjects

All experiments and procedures involving animals were performed in accordance with NIH guidelines and approved by the Institutional Animal Care and Use Committee of Northwestern University. A total of 26 adult C57BL/6J male mice (Jackson Laboratories stock #000664) were used in reported experiments and early pilot studies to establish methodology. All mice were individually housed under a 12-hour light/dark cycle during the course of the experiment. At the time of reported measurements, animals were 12-24 weeks old and weighed approximately 24-30 g. All animals were used for the first time in this experiment and were not previously exposed to pharmacological substances or altered diets.

### Surgical procedures

All surgeries were performed under isoflurane anesthesia (1-3%) in a stereotaxic frame. The surgical workflow consisted of two main procedures separated by a recovery and behavioral training period.

#### Headplate Implantation

First, mice were implanted with a custom 3D-printed plastic headplate. The headplate, which features an open-center design for cranial access, was affixed to the skull using dental cement (Metabond, Parkell). The position of bregma relative to marks on either side of the headplate was measured to facilitate the positioning of craniotomies during the subsequent surgery. The skull within the open window was then covered with a protective layer of dental cement. After a one-week recovery period, mice were placed on a water schedule (1-1.5 ml per day) and began behavioral training.

#### Craniotomies

After the completion of behavioral training, a second surgery was performed. The protective dental cement was removed to expose the skull, and four craniotomies (0.5-1 mm diameter) were drilled over the target regions. A platinum-iridium ground wire soldered to a connector pin was implanted in a separate craniotomy over the right frontal lobe and cemented to the headplate implant. Silicone elastomer (Kwik-Cast, World Precision Instruments) was used to cover the craniotomy after surgery.

### Probe insertion trajectory planning

Probe insertion trajectories were planned using a reference mouse brain atlas^117^ and the Allen Mouse Brain Common Coordinate Framework (CCFv3)^118^, aided by the “Neuropixels trajectory explorer software” (https://github.com/petersaj/neuropixels_trajectory_explorer). Targets in the left hemisphere included mPFC, M2, M1, MD, and GP. Targets in the right hemisphere were V1, SL and DCN. Table S1 lists the coordinates for the four probe trajectories (AP: antero-posterior; ML: mediolateral; DV: dorso-ventral).

**Table S1.** Stereotaxic coordinates for craniotomies and Neuropixels probe trajectories.

| Trajectory | Hemisphere | Target regions | Craniotomy coordinates (AP, ML) | Top electrode (AP, ML, DV) | Bottom electrode (AP, ML, DV) |
| --- | --- | --- | --- | --- | --- |
| 1 | Left | mPFC, M2 | 2.25, 1.04 | 2.35, 1.27, 0.38 | 1.92, 0.24, 4.06 |
| 2 | Left | M1, GP | -0.60, 1.11 | -0.60, 1.20, 0.69 | -0.60, 1.70, 4.50 |
| 3 | Left | MD | -1.70, 1.80 | -1.61, 1.57, 0.45 | -1.04, 0.08, 3.94 |
| 4 | Right | V1, SL, DCN | -3.91, -3.28 | -4.15, -3.20, 0.58 | -6.49, -2.34, 3.50 |

### Climbing apparatus

The behavioral apparatus, described in detail previously^67^, was housed inside a sound-attenuating chamber (H10-24A, Coulbourn). The core of the apparatus was a custom, 3D-printed cylindrical wheel featuring handholds positioned 12° apart. A ratchet mechanism permitted the wheel to rotate only in one direction, downward as the mouse climbed. The wheel’s absolute angular position was continuously tracked by a shaft encoder (A2-A-B-D-M-D, U.S. Digital). Experimental control utilized custom MATLAB (MathWorks) scripts and a DAQ card (PCI-e-6323, National Instruments).

### Behavioral training

Mice were initially acclimated to being handled by the experimenter, to head-fixation, and to the climbing apparatus according to previously described procedures^67^. Briefly, following acclimation to handling and head-fixation in front of the wheel, mice underwent several daily training sessions to associate rotating the wheel with a water reward. Initially, rewards were delivered manually for any wheel rotation. Subsequently, an automated protocol rewarded mice for performing progressively longer climbing bouts, with the reward amount for each bout adapted based on the mouse’s performance on recent bouts. This helped maintain motivation.

Once mice were proficient, they were transitioned to the behavioral protocol also used during recording sessions. This protocol consisted of two period types – “GO periods” and “NO-GO periods” – occurring alternately. So that mice could distinguish between periods, auditory (9 kHz pure tone; Murata Electronics 7BB-12-9) and visual (green led; Visual Communications Company CNX722C50005B) cues were presented during one of the two period types. Which period involved cues was varied across animals: for some mice, cues were active during the GO periods, while for others, they were active during the NO-GO periods. The duration of the GO and NO-GO periods was contingent on the mouse’s behavior. If the animal remained immobile, the durations of both GO and NO-GO periods were randomly selected from a continuous uniform distribution ranging from 2 to 4 seconds. If the mouse climbed, period duration was adjusted as follows:

- During a GO period, a climbing bout would terminate the period and trigger a reward.

- During a NO-GO period, a climbing bout would forestall the end of the period and reset the timer from the moment immobility is detected.

The present study focuses exclusively on the self-initiated climbing bouts, which occurred during NO-GO periods. A typical behavioral session lasted one hour.

### Video recording

Video tracking of the behavior was done using a high-resolution monochrome camera (acA2040-55um, Basler), fit with a 16-mm fixed focal length lens (33-304, Edmund Optics). The camera was positioned to the right of the head-fixed mouse and videos were acquired continuously throughout behavioral sessions. Recording was performed under near-infrared illumination at 100 frames per second with a resolution of 800 x 700 pixels.

To ensure precise temporal alignment, video acquisition was hardware-triggered. A 100 Hz TTL pulse train was generated by the NI-DAQ card, with each pulse triggering the acquisition of a single camera frame. This pulse train was recorded by the NI-DAQ system, providing a common temporal reference for the subsequent synchronization of the video frames with all other data streams, including electrophysiology and behavioral signals like wheel position.

We used DeepLabCut^68^ to track the positions of the mouse’s right hand and foot from videos. The resulting marker coordinates were then used to compute kinematic variables, such as limb velocity.

### Electrophysiological recording

Prior to each acute recording session, several preparatory steps were taken. To facilitate smooth penetration through the dura mater, the tips of the four Neuropixels 1.0 probes (IMEC) were sharpened using a grinder (EG-45, Narishige). A few minutes before insertion, the sharpened tips were coated with a fluorescent lipophilic dye (CM-DiI, Invitrogen)^119^ to mark the probe tracks for posthoc histological verification, according to the following protocol: https://www.protocols.io/view/painting-neuropixels-probes-and-other-silicon-prob-5jyl8neprl2w.

For the recording session, the mouse was head-fixed in the climbing apparatus and the silicone elastomer was removed from the craniotomies. The four Neuropixels, mounted on 3-axis micromanipulators (New Scale Technologies), were then slowly lowered to their target depths. The exposed skull was covered with silicone oil. Electrode voltages were acquired at 30 kHz using the Open Ephys acquisition software^120^. A 1 Hz TTL pulse train was used to align Open Ephys and Matlab recordings, including video.

### Spike sorting and firing rate estimation

Raw electrophysiological data was spike-sorted using Kilosort 3^121,122^. Only units classified as “good” single units by Kilosort were retained for further analysis. To estimate the instantaneous firing rate for each neuron, the binary spike trains (1 kHz) were convolved with a Gaussian kernel (σ = 10 ms)^123^. The resulting smoothed signals were then binned into 50-ms windows to compute the firing rate time series used in subsequent analyses. An exception was made for the analyses presented in Figures S3C and S3E: to capture dynamics at a faster timescale, these firing rate time series were derived by binning raw spike trains into 10-ms windows without prior Gaussian smoothing. Finally, we applied a selection criterion to focus on task-relevant neurons: only units with a mean firing rate of at least 0.5 Hz averaged across all periods from -1 s to -0.25 s relative to self-initiated climbing onset were included. These units are referred to as “qualifying units”.

### Histological procedure and brain registration

At the conclusion of the recording sessions, mice were deeply anesthetized and transcardially perfused with 3% paraformaldehyde (P6148, MilliporeSigma). Brains were extracted, post-fixed in paraformaldehyde, and subsequently sectioned into 60-µm thick coronal slices using a sliding freezing microtome (SM2000, Leica). The sections were then mounted on glass slides with a mounting medium containing DAPI (00-4959-52, Invitrogen). Whole-slide images were acquired using a slide scanner fluorescence microscope (VS120, Olympus). To verify the final position of probe tracks, the resulting images were registered to CCFv3^118^ using the open-source Python package HERBS^124^.

### Climbing onset detection

To precisely identify the onset of self-initiated climbing bouts, we used a two-stage method that combined wheel rotation and limb kinematics.

1. First, during behavioral sessions, climbing bouts were detected using the angular displacement of the wheel. The shaft encoder signal, acquired at 1 kHz, was smoothed, and a climbing bout was defined as a continuous rotation exceeding an angular displacement of 10°. Immobility, in this context, was defined as less than 0.25° of rotation over 400-ms windows. This step ensured that detected movements corresponded to genuine locomotion that displaced the wheel.
2. Second, this detection was refined offline using the kinematic data from the right hand and foot to determine the precise moment of climbing onset. The DeepLabCut marker coordinates were upsampled to 1 kHz via Akima spline interpolation^125^, and instantaneous velocity was calculated for each limb. A velocity threshold, empirically set to 0.1 pixel/ms to separate limb movements from baseline noise, was applied to classify time points as either mobile or immobile. Among periods of mobility, to distinguish genuine climbing from twitches, we applied the following criteria: 1) brief periods of mobility lasting less than 100 ms were classified as twitches, and 2) short pauses of immobility (< 400 ms) within a larger movement were reclassified as part of the ongoing movement. The onset of a potential climb was defined as the first moment a limb transitioned from an immobile to a mobile state that was not a twitch. For a potential climbing bout to be validated, the kinematics-defined onset had to be followed by a wheel-defined onset within 500 ms, ensuring the limb movement resulted in a productive wheel displacement exceeding 10°. Finally, the climbing bout had to be preceded by a continuous period of immobility of at least 1 second.

A single trial was defined as the continuous period of immobility preceding the onset of a climbing bout initiated during a NO-GO period.

### Quantification of anticipatory licking behavior

To quantify anticipatory licking behavior, we tracked individual licks during climbing from video recordings. First, using a custom MATLAB routine, we identified potential licks by detecting contrast changes within a region of interest between the water spout and the mouse’s mouth. Second, to eliminate false positives, which could occur when the mouse’s paw crossed the region of interest during climbing, every potential lick was manually curated by visualizing the corresponding frames.

We compared the anticipatory lick rate between GO and NO-GO periods by calculating the fraction of climbing bouts during which at least one lick occurred. To ensure an unbiased comparison between the two conditions, we enforced two criteria:

1) To prevent reward-triggered licking from contaminating the GO-period data, climbing bouts during GO periods were truncated immediately prior to the moment of bout completion, before the animal could perceive the reward or any reward-predicting cue. This truncation point corresponded either to the offset of the cues (for mice trained with active auditory and visual cues during GO periods) or to the moment of completion of a climbing bout (for mice trained with active cues during NO-GO periods, where GO period reward attainment is uncued prior to water delivery).

2) Because climbing bouts initiated during NO-GO periods are not rewarded, they tend to be longer than those during GO periods. To prevent this duration disparity from biasing the lick probability, we defined a session-specific “evaluation window” time-locked to climbing onset. Licking was only assessed within this window, and only climbing bouts (truncated as per criterion 1, if applicable) that met or exceeded this duration were included in the analysis. The length of this evaluation window was determined per session to resolve a trade-off: longer evaluation windows increase the cumulative probability of observing a lick, but simultaneously decrease the pool of eligible climbing bouts. To optimize this balance, we set the evaluation window to the median duration of all climbing bouts (truncated as per criterion 1, if applicable) in a given session. Consequently, our analysis retained half of the climbing bouts for each session, with licking evaluated over a period equal to this median duration.

### Cross-validated predictions of time-to-movement

To test whether the timing of self-initiated actions was predictable from neural activity on single trials, we trained a linear forecast model. We chose an elastic net linear regression model^76^ for its simplicity and interpretability. The model was trained to predict the time remaining until action onset (“time-to-movement”) at 50-ms time steps during trials. Trials, which had variable durations, were aligned to action onset. The input features for the model at each 50 ms time step consisted of the firing rates of all qualifying units, over the preceding five time steps (250 ms). We used a 5-fold crossvalidation scheme to train and test the forecast model. The regularization parameter for the elastic net was optimized for each fold using an internal 5-fold cross-validation loop performed exclusively on the training set. To assess the correlation of single-trial forecast predictions across brain regions (Figures 3D-3G), the same procedure was applied using qualifying units from individual regions.

To estimate chance-level predictions, we circularly permuted^126^ the time-to-movement relative to the firing rates for the training data 1,000 times for each cross-validation fold, and tested the resulting null models on the held-out test trials. We could thus compute a null distribution for each performance metric (mean absolute residuals and best-fit slopes) for each test trial. Actual performance metric values were compared against the null distributions to compute a *p*-value for each performance metric (Figures 2C-2E).

These calculations were repeated for 250-ms windows that began every 50 ms during a trial. This produced a sequence of *p*-values over time for each trial, where each *p*-value represented the forecast’s significance within that specific time window. We applied the Benjamini-Hochberg procedure^127^ to control the False Discovery Rate (FDR) across all time windows within a given trial. The onset of predictability for a trial was then defined as the first time point at which the FDR-corrected *p*-value fell below the significance threshold of 0.05.

### Control analysis to exclude “clock-like” activity

To test the alternative hypothesis that the neural activity used by our forecast model was merely tracking the passage of time from immobility onset, rather than reflecting a decision-related process, we performed a control analysis using synthetic “clock-like” activity (Figures S2A-S2C). We generated a synthetic dataset consisting of trials with variable durations, randomly drawn from a uniform distribution between 1 and 5 s. For each trial, we simulated a one-dimensional latent variable modeled as a linear ramp to which we added independent Gaussian noise. This signal mimics the readout of a neural population encoding the elapsed time since immobility onset. This synthetic dataset was then processed through the exact same 5-fold cross-validated elastic net forecast procedure used with the actual neuronal data. We then compared the relationships between forecast parameters (initial value, slope, final value) and trial duration in the synthetic data (Figures S2A-S2C) versus in the actual neural recordings (Figures 4A-4D).

### Random subsampling validation

We performed a random subsampling analysis to verify that the negative correlation observed between the forecast initial value and trial duration (Figure 4A) was a robust feature of our data, rather than being driven by a small subset of trials where the decision process started very early after immobility onset. We performed 10,000 iterations in which we randomly sampled 10% of the total trials without replacement. For each iteration, we computed the Pearson correlation coefficient and the associated *p*-value between the initial forecast values and the trial durations (Figure S4).

### Identification of forecasting neurons

To identify which individual neurons contributed significantly to time-to-movement predictions (Figures 3A-3C), we implemented a permutation-based feature importance analysis. The procedure was repeated 1,000 times; within each iteration, the following steps were performed for each of the five cross-validation folds:

1. For each recording session, an elastic net model was trained on training trials with all qualifying units.
2. On the held-out test data, we assessed the importance of each qualifying unit that had been assigned a non-zero coefficient by the model. To do this, we created a test set where the temporal information of a single neuron was destroyed by circularly permuting its activity time-series, while leaving the activity of all other neurons intact.
3. The original, already-trained model was used to predict time-to-movement from this partially permuted test data.
4. The effect of permuting a neuron was quantified by the change in mean absolute residual (MAR) compared to the MAR on the original test data. A neuron that improves the forecast should cause the MAR to increase when its activity is shuffled.

This process yielded a [1000 repetitions x 5 folds] matrix of MAR differences for each neuron. To determine statistical significance, we performed a non-parametric sign permutation test on this matrix. The null hypothesis of this test is that there is no consistent effect of shuffling a neuron’s activity, meaning the sign of the MAR difference is random across folds and repetitions. The observed statistic was compared against a null distribution generated by randomly flipping the signs of the differences in the matrix (10,000 permutations). Finally, the resulting *p*-values for all neurons were corrected for multiple comparisons using the Benjamini-Hochberg FDR procedure. A neuron was classified as a “forecasting neuron” if its FDR-corrected *p*-value was less than 0.05.

### Cross-correlation analyses

To quantify temporal offset between brain regions in their predictive activity, we performed crosscorrelation analyses. For each pair of brain regions, cross-correlograms were computed on each trial’s time-to-movement predictions (including data up to 250 ms before action onset). The empirical delay for a given pair of regions (Figures S3B and S3C) was defined as the time lag that yielded the maximum positive correlation within this trial-averaged cross-correlogram.

To assess whether the observed temporal delay between regions was statistically significantly different from zero, we employed a non-parametric bootstrap procedure. For each pair of regions, we generated 1,000 bootstrap samples. Each sample was constructed by randomly drawing *N* trials with replacement from the original pool of *N* trials. For every bootstrap iteration, we averaged the crosscorrelograms of the resampled trials and extracted the time lag corresponding to the new peak correlation. This procedure generated an empirical distribution of 1,000 expected lags, from which we defined the 95% confidence intervals of the true lag. Statistical significance was determined using a two-tailed test on the bootstrap lag distribution. Finally, to account for multiple testing across all regional pairs, raw p-values were adjusted using the Benjamini-Hochberg FDR procedure, with an alpha threshold set at 0.05.

### Drift-diffusion modelling

DDMs were fit to time-to-movement forecast traces derived from the entire neural population (Figure 5) and from individual regions (Figure 6). Separate models were fit for each recording session.

#### Model formulations and likelihood functions

Our modeling approach is aimed at describing the evolution of a decision variable *x* over time. It starts from an Ornstein-Uhlenbeck process^89,128^ and incrementally adds parameters to build towards the models reflected in Equations 1 and 2-5. All continuous-time stochastic differential equations (SDEs) were discretized using the Euler-Maruyama method, a standard numerical procedure for approximating the solution of SDEs^129^, with a time step of Δ*t* = 50 ms matching the bin size of firing rate time series.

Let *x*_*n*_ denote the value of the decision variable at time step *n*, corresponding to time *t*_*n*_ = *n*Δ*t*. The process begins at immobility onset (*n* = 0, *t* = 0) with a latent initial state *x*_0_. The first observed data point in our forecast traces corresponds to *t* = 250 ms after immobility onset, which is time step *n* = 5. Thus, a single trial trajectory is observed as the sequence ***x***_***obs***_ = [*x*_5_, *x*_6_, …, *x*_*N*_]. We explicitly modeled the unobserved values of *x* between immobility onset (*x*_0_) and the first observation (*x*_5_). The process was initialized at *n* = 0 as 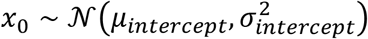. The mean and variance were recursively propagated forward from the initial state *x*_0_ to the first observed time step *x*_5_ (*n* = 5, *t* = 250 ms) according to each model’s dynamics. This yielded 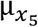 and 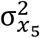, the conditional moments for the first observed point 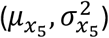 against which the model likelihood for *x* was computed. For the remainder of the trajectories (*t* > 250 ms), the likelihood of Models 1 to 3 (see below) was evaluated step-by-step by computing the probability of each new observation given the model dynamics propagated from the previous empirical data point.

To understand the derivation of the likelihood functions below, consider a general SDE of the form *dx*_*t*_ = *A*(*x*_*t*_, *t*)*dt* + *cdW*_*t*_. The Euler-Maruyama discretization yields the recursive update *x*_*i*+1_ = *x*_*i*_ + *A*(*x*_*i*_, *t*_*i*_ )Δ*t* + *c*η_*i*_, where the noise increment η_*i*_ is drawn from a normal distribution 𝒩(0, Δ*t*). Consequently, the probability of transitioning to the next state is normally distributed: *P*(*x*_*i*+1_|*x*_*i*_ ) = 𝒩(*x*_*i*+1_; *x*_*i*_ + *A*(*x*_*i*_, *t*_*i*_)Δ*t, c*^2^Δ*t*). Because standard white noise has independent increments, the process satisfies the Markov property: the state at *t*_*i*+1_ will depend only on the state at *t*_*i*_. The joint likelihood of a full trajectory is therefore the product of the initial state probability and all subsequent step-by-step transition probabilities, which translates to a sum of log-normal densities when computing the log-likelihood.

This step-by-step multiplication is only valid if the noise increments are independent. In Model 4, where noise is colored (fractional Gaussian noise), the increments are autocorrelated, violating the Markov property. In that case, the probability of transitioning to *x*_*i*+1_ depends not only on *x*_*i*_ but on the entire history of the noise, necessitating a multivariate normal evaluation of the entire trajectory rather than a product of local transitions.

#### Model 1: Ornstein-Uhlenbeck process (OUP; 4 parameters)

This baseline model represents a leaky accumulator with no drift term. Its parameters are parameters are θ_*Model*1_ = [*c, k*, μ_*intercept*_, σ_*intercept*_]. The dynamics are governed by the SDE:

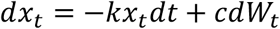

where *dW*_*i*_ represents the increments of a standard Wiener process (white noise).

Its discretized form is:

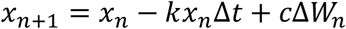

The log-likelihood for a single trajectory *x* is given by:

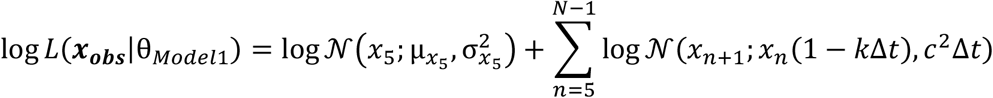

where 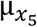 and 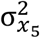 are obtained by propagating the initial state parameters (μ_*intercept*_, σ_*intercept*_ ) through the leak and noise accumulation for the first five time steps (*t* = 0 → 250 ms).

#### Model 2: Constant drift model (5 parameters)

This model adds a constant drift rate, *I*, to the OUP, with parameters θ_*Model*2_ = [*c, k*, μ_*intercept*_, σ_*intercept*_, *I*.] Its SDE is:

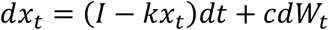

Its discretized form is:

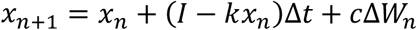

The log-likelihood for a single trajectory is given by:

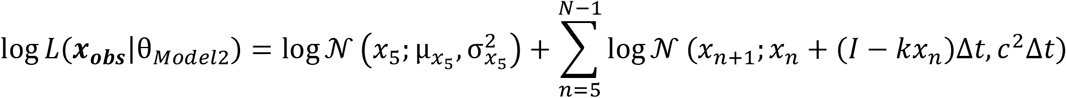

where 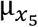 and 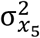 are obtained by propagating the initial state parameters (μ_*intercept*_, σ _*intercept*_) through the model for the first five time steps. The contribution of a constant drift was tested by comparing the likelihoods of Models 1 and 2.

#### Model 3: Full model with white noise (7 parameters)

This is our primary model described in the main text (Equations 2-5), which incorporates trial-to-trial variability in the initial drift rate and a time-dependent drift rate *I*(*t*). Model parameters are θ_*Model*3_ = [*c, k*, α, μ_*intercept*_, σ_*intercept*_, μ_*drift*_, σ_*drift*_ ].

Its SDE is:

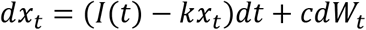

To account for trial-to-trial variability, the initial drift magnitude *I*_0_ is treated as a latent random variable following a lognormal distribution log 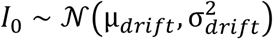. This distribution was chosen to constrain the drift rate to positive values. The drift can evolve monotonically over time according to either an exponential function *I*(*t*) = *I*_0_*e*^α*t*^ or a linear function *I*(*t*) = *I*_0_ + α*t*.

The discretized form is:

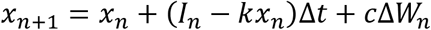

with *I*_*n*_ = *I*_0_*e*^α(*n*Δ*t*)^ (exponential) or *I*_*n*_ = *I*_0_ + α(*n*Δ*t*) (linear).

The total log-likelihood across all trials **X**_***all***_ is computed by marginalizing over the initial drift magnitude *I*_0_:

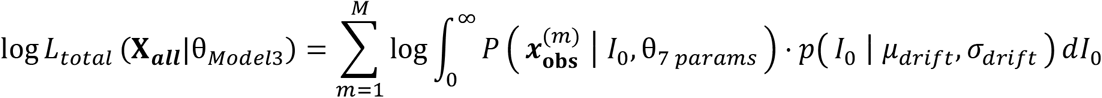

where *M* denotes the total number of trials and 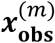 represents the observed trajectory of the *n*-th trial. *P*( ***x*** ∣ … ) denotes the conditional likelihood of a trajectory, while *p*( *I*_0_ ∣ … ) represents the probability density function of the initial drift magnitude.

The conditional log-likelihood of a single trajectory given a specific *I*_0_ is:

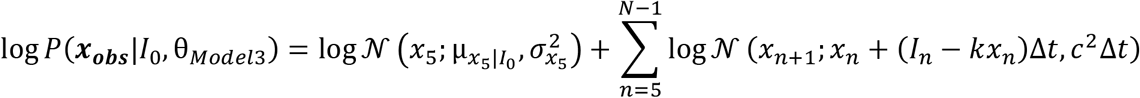

Here, 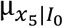 and 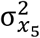 are derived by propagating the initial state parameters (μ_*intercept*_, σ_*intercept*_ ) through the model for the first five time steps. The contribution of the initial state variability (σ_*intercept*_), leak rate (*k*) and urgency (α) parameters was tested by comparing this full model to nested versions where either *σ*_*intercept*_, *k*, or α was fixed to zero.

#### Model 4: Full model with colored noise (8 parameters)

To test for temporal autocorrelations in the noise, we extended the full model by replacing the standard white noise process with fractional Gaussian noise, characterized by a spectral exponent parameter β. This model’s parameters are θ_*Model*4_ = [*c, k*, α, μ_*intercept*_, σ_*intercept*_, μ_*drift*_, σ_*drift*_, β.] The corresponding SDE is:

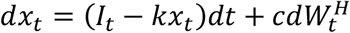

where 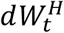 represents the increments of a fractional Brownian motion. The parameter β is defined as the spectral exponent of the noise power spectrum (*S*(*f*) ∝ 1/*f*^β^ where *S*(*f*) denotes the power spectral density at frequency *f*), which relates to the Hurst exponent by *H* = (β + 1)/2.

Unlike the white noise models where transitions are independent, the presence of colored noise introduces correlations between time points within a trajectory. Consequently, the conditional likelihood of a full observed trajectory ***x***_***obs***_ given a drift *I*_0_ is evaluated using a single multivariate normal distribution:

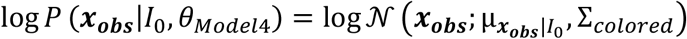

Any observed trajectory remains defined from the 5th step onward (i.e., ***x***_***obs***_ = [*x*_5_, *x*_6_, …, *x*_*N*_ ]). The mean vector 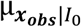 is constructed by propagating the initial state mean μ_*intercept*_ and the trialspecific drift *I*_0_ through the deterministic dynamics of the model from immobility onset (*n* = 0) up to the observed steps.

To evaluate the likelihood, we first derived a relation for the covariance matrix Σ_*colored*_, which captures the dependency structure resulting from the discrete leaky accumulation of colored noise. We analytically derived its elements from the recursive equation for the model. First, since the noise is additive (i.e., the diffusion amplitude *c* is independent of the state *x*_*t*_), the continuous-time SDE can be integrated pathwise and straightforwardly discretized using an Euler-Maruyama scheme^130,131^. With a time step Δ*t*, this yields:

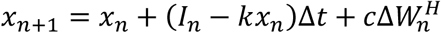

Defining the discrete-time leak decay factor as ϕ = 1 − *k*Δ*t* and the discrete noise increment as 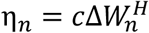, the recursive update equation becomes:

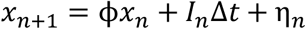

Unrolling this recursive equation from immobility onset (*n* = 0) to an arbitrary time step *n* yields *x*_*n*_ as the sum of the leak-attenuated initial state, the drift, and the accumulated noise:

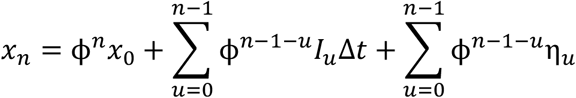

With this decomposition of *x*_*n*_, we can now derive the exact dependency structure between any two time points. Specifically, the elements of the covariance matrix Σ_*colored*_ (*n*_*t*_, *n*_*b*_) represent the covariance between the states at any two time steps *n*_*t*_ and *n*_*b*_ (where *n*_*t*_, *n*_*b*_ ≥ 5, corresponding to the observed time steps) conditioned on a specific trial drift *I*_0_:

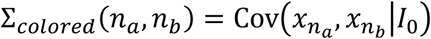

Because the drift component is deterministic for a trial-specific initial value *I*_0_, it strictly shifts the mean of the trajectory 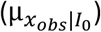 without contributing to the covariance of the fluctuations. Consequently, the covariance structure depends only on the attenuated initial state variability and the history of the colored noise increments. Furthermore, the initial state 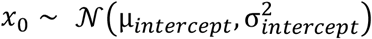 is statistically independent of the subsequent within-trial noise increments η. Applying the covariance operator 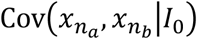 thus simplifies to the sum of two independent covariance terms:

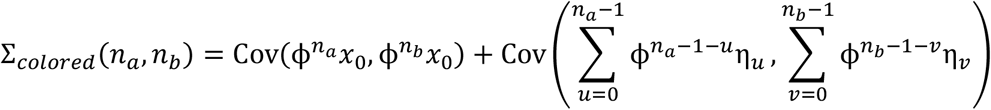

For the first term, factoring out the scalar multipliers, 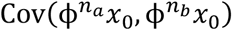 becomes 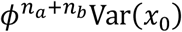, which equals 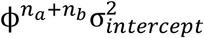. For the second term, the covariance of the sums expands into a double summation of the covariance between individual noise increments:

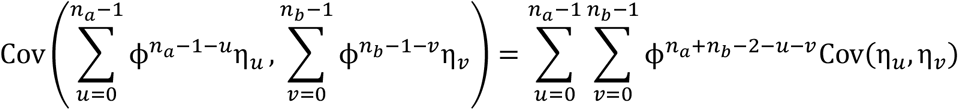

In discrete time, the covariance between two fractional Gaussian noise increments separated by a lag τ = |*u* − *v*| is proportional to the autocorrelation function ρ, such that Cov(η_*u*_, η_*v*_) = *c*^2^Δ*t*ρ(|*u* − *v*|)^132^.

Substituting these relationships yields:

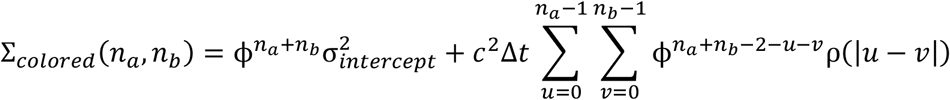

The first term represents the propagated variance from the initial state, while the double summation represents the convolution of the noise autocorrelation with the system’s impulse response (that is, the discrete-time leak decay). Analytically, the autocorrelation function of the fractional Gaussian noise is given by^132^:

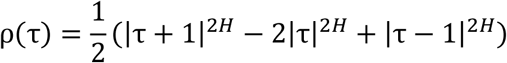

where τ = |*u* − *v*| is the lag between noise increments.

Finally, the total log-likelihood across all trials ***X***_***all***_ is computed by integrating this multivariate conditional likelihood over the lognormal prior distribution of *I*_0_:

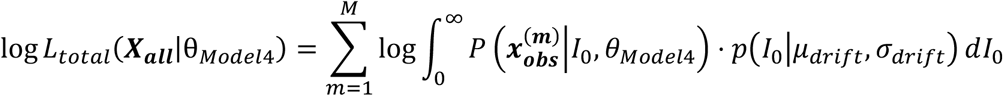

The parameter β was constrained to be between -1 and +1 to ensure the stationarity of the noise process^132^. Its contribution was tested by comparing the likelihoods of Models 3 and 4.

#### Data fit by the DDM

DDM parameters were estimated by fitting forecast traces derived from held-out test trials, spanning from their beginning up to 250 ms before action onset. The forecast traces were fit from their initial value (250 ms after immobility onset) based on our finding that this initial value is correlated with trial duration (Figures 4A and S4), and that predictive information typically emerges within the first 500 ms for most trials (Figure 2H), consistent with a decision-making process starting soon after immobility onset. The end point (250 ms before action onset) was chosen to isolate the decision-making process and avoid contamination from potential post-decision, motor execution-related activity. This choice should be conservative: in humans, the point of no return beyond which a self-initiated action can no longer be aborted occurs approximately 200 ms before action onset^15,78^. This delay is likely shorter in mice, given the much lower limb inertia.

As a pre-processing step, because our DDM formulation assumes a positive drift rate and leak rate, the forecast traces were first made non-negative by subtracting from all forecast traces the minimum value observed across all traces within that session.

#### Optimization and parameter fitting

Model parameters were fit by minimizing the negative log-likelihood of the data. We used a two-stage global optimization procedure to avoid local minima. First, a genetic algorithm^133,134^ was run five times with different random seeds to broadly explore the parameter space (function *ga* in MATLAB). The best-fit parameters from these runs were then used as the starting point for an interior-point local optimizer^135,136^ to refine the minimal negative likelihood estimate (function *fmincon* in MATLAB). Parameters such as leak rate, noise amplitude, and distribution widths (*k, c*, σ_*intercept*_, σ_*drift*_) were constrained to be positive, while the drift’s temporal rate α was allowed to be positive or negative.

#### Statistical procedure for model selection

The improvement in model fit for each parameter was assessed using a non-parametric bootstrap procedure^137^. We performed 1,000 bootstrap iterations. For each iteration, we constructed a training set by sampling *N* trials with replacement from the trials for the given session, where *N* is the total number of trials for that session. The remaining trials that were not selected in the training set served as the held-out test set. We fit full and nested models on the training set and evaluated their goodness-of-fit on the test set by computing the total log-likelihood for all held-out trials. To ensure comparability between bootstrap iterations and sessions, the total log-likelihood was normalized by the number of data points in the given test set. For each iteration, we computed the log-likelihood gain (Δ*LL*) as the difference between the full and nested models; these values are reported in Figures 5, 6, and S6. A positive Δ*LL* thus indicates that the additional parameter improved the model fit. Statistical significance was determined via an empirical *p*-value, defined as the fraction of bootstrap iterations where the full model failed to outperform the nested model (Δ*LL* ≤ 0). A parameter was considered to significantly improve the model if *p* < 0.05. This model selection approach was validated using parameter recovery on simulated data (Figure S5).

#### Derivation of drift rate moments

In our DDM variant, the initial drift rate *I*_0_ is drawn from a log-normal distribution for each trial. The parameters directly estimated by our fitting procedure, μ_*drift*_ and σ_*drift*_, correspond to the mean and standard deviation of the underlying normal distribution for *log*(*I*_0_) (Figures 5 and 6). For visualization purposes (Figures S6A, S6B and S6G), we also calculated the arithmetic mean (*driftmean*) and standard deviation (*driftstd*) of the log-normal distribution itself using the following equations:

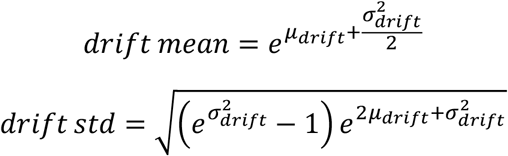

#### Testing the necessity of within-trial noise via ablation analysis

To assess whether the noise accumulation is necessary for reaching the activity threshold, we performed an ablation analysis on the diffusion term. Using the best-fit parameters obtained with the 7-parameter model (Equations 2-5), we simulated decision variable trajectories under two conditions: with the diffusion term (*c* set to its best-fit value) and without it (*c* = 0). To ensure the simulations were realistically constrained by observations, the threshold beyond which each simulated trajectory ended was drawn from a normal distribution fit to the actual forecast traces at 250 ms prior to climbing onset. This time point corresponds to the end of the DDM fitting window and was chosen to define a realistic activity threshold. To guarantee an unbiased comparison between the duration of simulations and actual trials, we discarded simulations shorter than the minimum permitted trial duration fit by the DDMs. The duration of the forecast traces fit by the DDMs equals the total trial (immobility) duration minus 0.45 s. This deduction accounts for a 0.2 s initial offset because the forecast model uses five time steps, and a 0.25 s truncation prior to climbing onset. Since the minimum trial duration is 1 s, the minimum fitted trial duration is 0.55 s. Accordingly, the duration of simulated sequences was calculated starting at 250 ms after their onset, and only simulated sequences lasting at least 0.55 s were retained for this comparison.

#### Assessment of distinct variability sources via dynamical constraints

To test that trial-to-trial variability in the decision process’s initial state (*σ*_*intercept*_) and the within-trial continuous noise arise from distinct sources of variability, we assessed whether the initial variance (*σ*_*intercept*_) could be explained by the noise accumulation during the period preceding immobility onset. For a process governed by leak rate *k* and noise amplitude *c* (i.e., an Ornstein-Uhlenbeck process), the variance derivative is 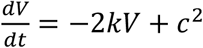, which yields the time-dependent solution 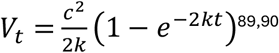. This function implies a theoretical upper bound, 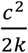 which represents the maximum variability the noise source can sustain against the leak, even if allowed to accumulate over an infinite duration. We compared the fitted initial variance 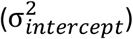 obtained from our 7-parameter model to this theoretical maximum calculated using the best-fit *c* and *k* parameters for each session (Figure S6C). Our finding of 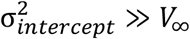 indicates that the initial variability exceeds the variability that could be attributed to noise accumulation.

## Acknowledgments

We are grateful to Daniel Dombeck, Christian Ethier, John-Dylan Haynes, Yevgenia Kozorovitskiy, Uri Maoz and Aaron Schurger for helpful discussions and feedback on an earlier version of the manuscript; to Carlos Brody for a helpful discussion; and to Daniel Greenberg for assistance with limb tracking. The computational analyses for this work were largely performed on Quest, Northwestern University’s high-performance computing cluster. This research was supported by a Searle Scholar Award, a Sloan Research Fellowship, a Simons Collaboration on the Global Brain Pilot Award and NIH grant DP2 NS120847 awarded to A.M.; a NIH grant R00 NS119787 awarded to J.G.; grants from the NSF (DMS-2235451) and Simons Foundation (MPS-NITMB-00005320) to the NSF-Simons National Institute for Theory and Mathematics in Biology.

